# Longitudinally cultured precision cut lung slices as a predictive *ex vivo* model for lung cancer

**DOI:** 10.1101/2024.08.23.609442

**Authors:** Ara Sargsian, Hermon Girmatsion, Vincent Lenders, Bart Ghesquière, Camila Takeno Cologna, Stefaan J. Soenen, Bella B. Manshian

## Abstract

Exploring alternatives to *in vivo* models, this study validates Precision Cut Lung Slices (PCLS) as a viable *ex vivo* platform for lung cancer research. We established the prolonged viability and structural preservation of PCLS, essential for accurate drug response studies. Using Paclitaxel as a benchmark drug and a therapeutically promising silver nanoparticles in combination with immunotherapy, we conducted a pioneering comparative analysis of its therapeutic effects on PCLS against traditional *in vivo* models. Results revealed that PCLS closely mimics *in vivo* responses, demonstrating comparable drug efficacy in tumor growth inhibition. This direct comparison not only confirms the utility of PCLS in simulating real-world outcomes but also emphasizes its potential in reducing animal testing. By providing a reliable, ethical, and efficient alternative for lung cancer studies, PCLS could significantly enhance preclinical research and drug development, marking a critical step towards more humane and representative scientific investigations.

## Introduction

The development of reliable disease models is keystone of biomedical research, representing an ongoing challenge in accurately replicating the intricate *in vivo* environment. Despite continuous advancements, the journey towards fully embodying the complexities of biological systems in a research setting remains a nuanced and evolving endeavor. This exploration seeks to address the multifaceted nature of current modeling techniques and their implications in the broader context of scientific inquiry and therapeutic development.

### From 2D to 3D Cultures: A Shift in Paradigm

Since their inception in the early 1900s, 2D cell cultures have been foundational in cellular and molecular biology, offering a controlled and manageable environment for studying cellular behavior, genetics, and biochemistry ^1,2^. The simplicity and ease of handling of 2D cultures have facilitated numerous breakthroughs in understanding disease mechanisms and screening potential drug therapies ^3^.

However, the flat and uniform nature of 2D cultures imposes significant limitations. Cells grown on flat surfaces lack the three-dimensional interactions and architecture inherent to actual tissues ^3^. This absence results in altered cell morphology, proliferation, and gene expression, leading to a physiological relevance gap ^4^. The absence of tissue-specific architecture and reduced cell-cell, and cell-matrix interactions in 2D systems oversimplifies the complex *in vivo* environment, often leading to results that do not translate effectively into actual biological processes ^3^.

In response to these limitations, 3D cultures have emerged as a more physiologically relevant platform. These systems allow cells to grow in all directions, better simulating the natural growth patterns, cellular interactions, and microenvironments of actual tissues ^5^. Organoids, spheroids, and other forms of 3D cultures have shown promise in providing a more accurate representation of the *in vivo* environment, thereby enhancing the predictive accuracy of preclinical studies ^4^. They offer a more dynamic and realistic model for studying tissue development, disease progression, and drug efficacy, as demonstrated by studies on host-pathogen interactions in lung slice cultures. For example, in studies involving Jaagsiekte sheep retrovirus, researchers observed dynamic viral replication over multiple time points. Similarly, chicken lung models maintained viability and functionality for over 40 days, allowing for real-time observation of immune responses and pathogen interactions ^4,6,7^. Additionally, murine precision-cut liver slices (PCLS) provide a dynamic tool for studying the tumor microenvironment and cell signaling *ex vivo*, including assays on cell adhesion to liver tissue, essential for understanding metastasis ^8^.

Despite their advancements, 3D cultures are not without limitations. While they provide a better approximation of tissue architecture and cellular interactions, they still require the artificial manipulation of cells and reliance on scaffold matrices, which may not fully mimic the extracellular environment found in living tissues ^9^. Moreover, these systems often lack the complete cellular diversity and complexity of actual tissue ^10^. Moreover, these systems often lack the complete cellular diversity and complexity of actual tissue. Healthy organoids are mainly made from iPSC-derived pluripotent stem cells. This results in a fabrication which, on the one hand, can result in large variability due to the efficacy of differentiation ^11^. Alternatively, it can also be limiting as the organoids would all be derived from the same cell source and thus do not represent the interindividual heterogeneity in tissue composition and architecture, whether healthy or diseased ^12,13^. Addressing these issues requires advanced co-culture systems and 3D culture techniques that incorporate multiple cell types and better mimic the *in vivo* environment to improve physiological relevance and reproducibility.

### The Translational Hurdle: From Animal Models to Clinical Trials in Biomedical Research

The evolution from 2D to 3D cultures marked a significant leap in accurately simulating the *in vivo* environment and tissue organization. Yet, even the most sophisticated *in vitro* models, like organoids and spheroids, cannot fully recapitulate the intricate biological processes that occur within living organisms. This is where genetically modified animal models have been pivotal, serving as a bridge to fill the crucial gap left by the 2- and 3D cultures, particularly in simulating human-specific disease mechanisms. However, the advent of advanced *ex vivo* models like PCTS (Patient-derived Tumor Slices) offers promising alternatives. These models can reduce the number of animals needed and bring studies closer to patient-specific conditions, especially in cancer research. With the FDA no longer requiring animal testing for drug approval, PCTS and similar technologies are poised to become valuable tools in preclinical research ^14^.

Genetically modified animal models, particularly those so-called “humanized” rodents that carry human genes, cells, tissues, or organs, have been instrumental in cancer research ^15–17^. These humanized models offer a more systemic context, allowing for the study of tumor growth, metastasis, and drug response within a whole-organism environment ^16,17^. This universal approach is vital as it encompasses not just the tumor cells but also the tumor microenvironment, including stromal cells, immune cells, and the vasculature, all of which play critical roles in cancer progression, treatment response, and novel therapeutics development ^18–20^.

However, while these models bring us closer to mimicking the complexity of human diseases, they are not without their challenges. One significant hurdle is the inter animal variability, which can lead to discrepancies in drug efficacy and toxicity. Moreover, the high cost, ethical considerations, and time required for developing and studying genetically modified animal models are substantial ^4,17,21^. Furthermore, given the mere 8% translatability of animal models to clinical trials, there’s a pressing need for advanced *ex vivo* models in biomedical research ^22^

The transition from 2D to 3D cultures, coupled with the use of transgenic animal models, has significantly advanced biomedical research, providing deeper insights into cellular behavior and tissue structure. Yet, the pursuit to emulate the complexities of human tissues persists, propelling researchers to explore more refined models and technologies, including Precision-cut Tissue Slices (PCTS).

### Bridging the Gap: Precision-cut Tissue Slices (PCTS)

PCTS represents a cutting-edge methodology, offering slices of live tissue that maintain the architectural, functional, and cellular complexity of the original organ. PCTSs are clearly defined tissue sections of uniform diameter and thickness, generated from a single organ, either human or animal. One of the distinguishing features of PCTS lies in its ability to preserve the spatial distribution of different cell populations within the original tissue ^23^. PCTS provides a unique platform that maintains the architectural, functional, and cellular complexity of the original tissue and functions as a valid *ex vivo* platform for respiratory diseases, viral and bacterial infection, rapid drug screening, immunotherapy, and drug response ^23–30^. PCTS has particularly been employed to understand and study alveologenesis, alcoholic liver injury, and antifibrotic therapies for fibrosis and cirrhosis ^24,31,32^.

Additionally, in alignment with the 3Rs principle (Refinement, Reduction, and Replacement), increasing efforts, including the adoption of precision-cut tissue slices, have been made to reduce the number of laboratory animals, thereby minimizing ethical concerns, and enhancing the relevance of research findings to human health ^33^. Therefore, PCTS are essential *ex vivo* model that captures the intricacies of the *in vivo* tumor microenvironment with high accuracy, offering distinct advantages over traditional organotypic models such as spheroids and organoids ^34^. In stark contrast to organoids, where enzymatic reactions, during generation, may disrupt the natural arrangement of cells and compromise the cell-cell and cell-matrix interactions ^23^. PCTSs are generated without the influence of enzymatic reactions, exhibiting a high degree of similarity to the original tumor, making them a valuable tool for researchers aiming to study the native behavior of cells in their microenvironment ^35,36^.

The utility of PCTS extends beyond traditional drug screening applications, particularly in the context of personalized medicine ^36,37^. With continuous advancements in combination treatments and the emergence of novel anti-cancer drugs, PCTS presents itself as an efficient platform for screening drugs and therapies tailored to individual patient profiles ^25,34,37^. Through cultivating PCTS from patient-derived biopsies, immediate access is gained to a representative *ex vivo* model without the need for extensive processing ensuring that the results obtained are more reflective of the *in vivo* scenario ^35^. While organoids and patient-derived xenografts (PDX) are often considered in similar contexts, the generation of these models is frequently marred by complexity, high cost, and labor-intensive procedures. Patient-derived xenografts, in particular, suffer from a low success rate, limiting their reliability as a consistent experimental model ^22^. Organoids, although encompassing various cell types found in patient tissues, fall short in replicating the spatial distribution and complex cell-cell and cell-matrix interactions present in the native tumor microenvironment ^23,35^. In particular, non-small cell lung cancer organoids were reported to have a success rate of only 17%, a limitation primarily attributed to the overgrowth of cancer cells by normal, healthy cells, as detailed in the findings of Dijkstra *et al*. ^38^.

Transitioning the focus to PCTSs, the commonly encountered challenges with organoids and PDX models can be addressed effectively, particularly in replicating complex biological environments. PCTS stands out as a potential solution, offering a more accurate *ex vivo* representation for *in vivo* experiments.

However, despite the successful generation of these models, an in-depth comparison with their *in vivo* equivalents is crucial to validate their efficacy. Recognizing this gap, we have generated PCTSs from an orthotopic mouse cancer model and thoroughly compared the *ex vivo* model with the equivalent *in vivo* model. This comparison was performed by employing *in vivo* imaging system (IVIS) screening following paclitaxel administration, ensuring a comprehensive analysis. Additionally, in our pursuit to enhance the utility of PCTS, we propose a more optimal freezing protocol aimed at preserving long-term tissue viability, further bridging the gap between *ex vivo* and *in vivo* methodologies.

In the subsequent sections, we aim to explore the potential of precision-cut tissue slices (PCTS) as a highly effective *ex vivo* model, building on their established use in various pathologies. This investigation will focus on assessing the strengths and applications of PCTS specifically in the realms of cancer research, drug development, and drug screening. By doing so, we seek to determine whether PCTS can provide a versatile and relevant platform for these critical areas, thereby offering novel insights and advancements in *ex vivo* modeling.

## Materials and Methods

### Preparation of lung tissue slices

Precision cut lung slices (PCLS) were prepared from healthy BALB/c mice lungs. After euthanasia and dissection of the animal, a small cut was made in the trachea, and the lung was injected with 1 - 2 mL of 2 % low melting point agarose (VWR, Belgium) in sterile Phosphate Buffered Saline (PBS; Gibco, ThermoFisher Scientific, Belgium) with a syringe, followed by putting the lung on ice for quick gelation of the agarose. This technique inflates the collapsed lung, restoring morphological structures while preserving the viability of the tissue. Additionally, agarose gel contributes to the overall structural strength of the sample, making it suitable for PCLS preparation. Cylindrical tissue cores of 5 mm were punched. PCLS (250 μm thick) were obtained using the Krumdieck tissue slicer (coring tool and slicer, Alabama Research and Development, Munford AL, USA) in ice-cold sterile PBS. The arm speed was set on the lowest speed and the blade speed was set on the second lowest speed. The slices were placed onto 24-well plates in different culture media for 1h in an incubator. The media were changed after 1 h and then every 2 days. The slices were incubated at 37 °C and 5% CO2 in 24-well plates.

### Culture media

Dulbecco’s Modified Eagle Medium/Nutrient Mixture F-12 (DMEM/F-12; Gibco, ThermoFisher Scientific, Belgium) supplemented with 0%, 1%, or 10% fetal bovine serum (FBS; Gibco, ThermoFisher Scientific, Belgium) 1% penicillin/streptomycin (Corning, Belgium) and 1% Amphotericin B (AK scientific, USA).

### PCLS viability (ATP / AlamarBlue HS / LDH)

The viability of PCLS was evaluated using various viability assays, including ATP, AlamarBlue HS, and -LDH. To measure the intra- and extracellular ATP levels, the CellTiter-Glo® 3D Cell Viability Assay (Promega, UK) was utilized according to the manufacturer’s instructions. The plate was incubated at room temperature for 30 minutes, and the luminescence values (open filter) were read using a microplate reader (SpectraMax iD3, Molecular devices). To assess the reducing capability of the viable cells in the PCLS, the slices were exposed to AlamarBlue HS (ThermoFisher Scientific, Belgium) according to the manufacturer’s instructions and incubated for 1 hour and 30 minutes at 37°C and 5% CO2 in 24-well plates. The fluorescent values (ex/em: 560/590) were then read using the same microplate reader.

The extracellular LDH was measured using Lactate dehydrogenase (LDH) assay kit (fluorometric) (abcam, UK) according to the manufacturer’s instructions. The supernatant was incubated for 10 min at RT. The fluorescent values (ex/em: 535/587) were then read using the same microplate reader.

### Glycolysis assay

To analyze the effect of long-time cultivation on glycolysis ability in PCLS, extracellular acidification rate (ECAR) was investigated using XF-24 cell culture assay (Seahorse Bioscience, North Bill-erica, MA, USA) according to the manufacturer’s procedure.

PCLS are typically cultured in glycolysis stress test medium without glucose, and their ECAR is assessed. To initiate the test, a saturating concentration of glucose is administered as the first injection. The cells take up and metabolize the glucose through the glycolytic pathway, resulting in the production of ATP and lactate. This process leads to the rapid increase of ECAR. The rate of glycolysis under basal conditions is represented by the glucose-induced response.

In the second injection, oligomycin, an ATP synthase inhibitor, is administered. Oligomycin inhibits mitochondrial ATP production, thereby shifting the energy production to glycolysis, and causing an increase in ECAR that reveals the maximum glycolytic capacity of the cells. The final injection is 2-DG, a glucose analog, which inhibits glycolysis by binding competitively with glucose hexokinase, the first enzyme in the glycolytic pathway. The subsequent decrease in ECAR confirms that the ECAR produced during the experiment is due to glycolysis.

The difference between the Glycolytic Capacity and the Glycolysis rate defines the Glycolytic Reserve. ECAR before the glucose injection is referred to as non-glycolytic acidification, which is caused by processes in the cell other than glycolysis.

Briefly, PCLS were prepared and cultured to get different time points (day 0, day 4, and day 8, n = 6 for each timepoint) on the day of assay. Drug compounds related to glycolysis ability (Glucose: 10 mM, Oligomycin: 1.0 μM, 2-DG: 50 mM) were mixed and added to the pH 7.4 culture medium. ECAR was analyzed using an XF analyzer (Agilent, Belgium) according to the manufacturer guideline.

### Gene expression assay

PCLS (healthy and cancerous, n =3) were put, on different time points, in RNA later (in RNAse free vial) overnight at 4°C and stored at -80 °C without RNA later. On the day of processing, the tissues were put in an empty 2ml RNAse-free vial with a stainless steel bead (Qiagen, Germany) and homogenized using a tissue homogenizer (frequency: 25, time: 30 sec) (TissueLyser II, Qiagen, Germany). RNA was isolated using a RNeasy mini kit (Qiagen, Germany) according to the manufacturer’s instructions. The quality of the isolated RNA was verified using nanodrop (ND-1000 Spectrophotometer, Isogen Life Science, Netherlands). cDNA was prepared using iScript Select cDNA synthesis kit (Bio-Rad, Belgium) according to the manufacturer’s instructions. The obtained cDNA was used for RT-qPCR (StepOnePlus, Thermo Scientific) to detect the expression of the following genes:

All the primers were purchased from integrated DNA technologies (Belgium, Leuven). PCR conditions were 95 °C for 2 min followed by 40 cycles of 95 °C for 5 s and 60 °C for 25 s.

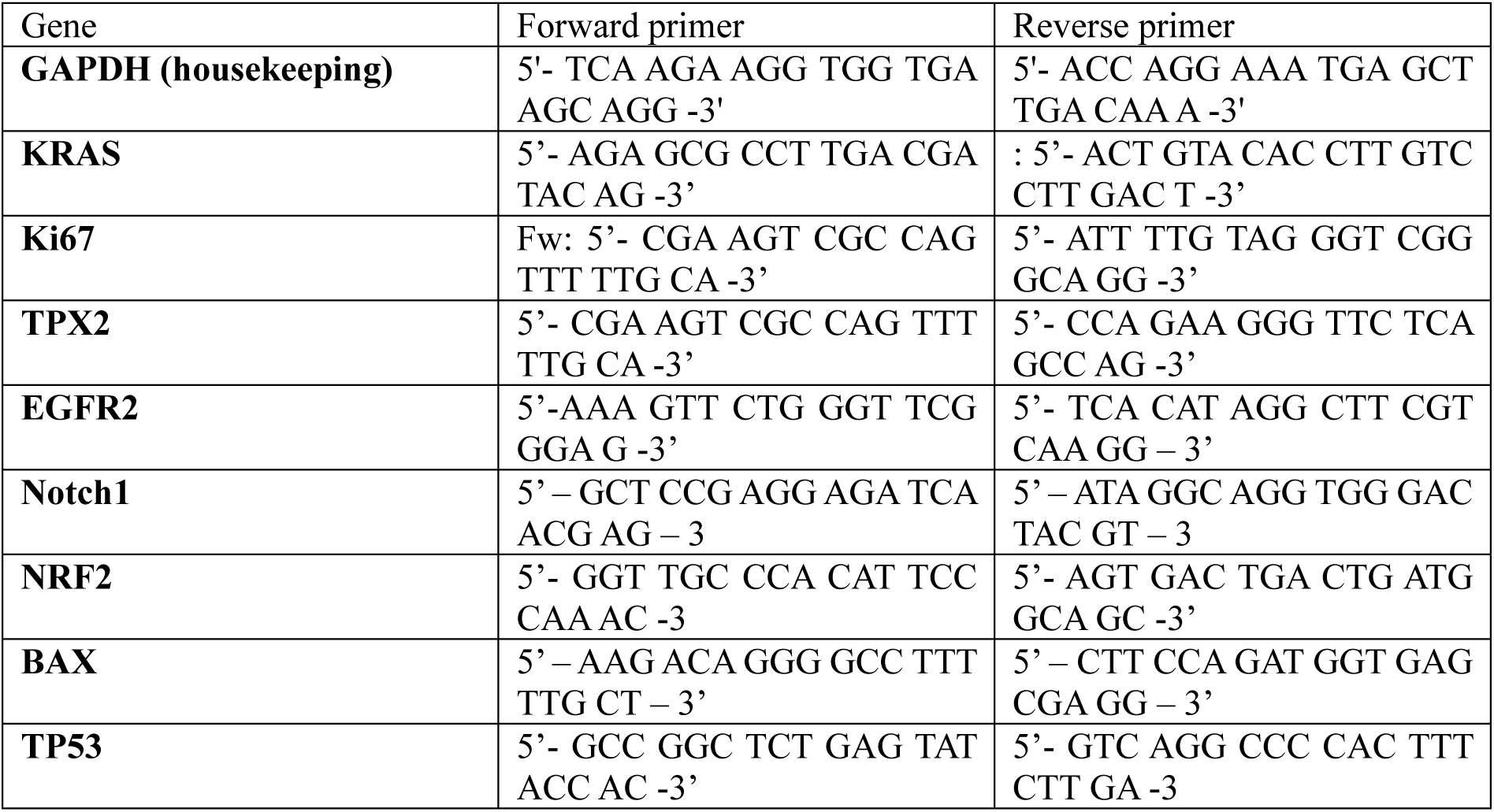
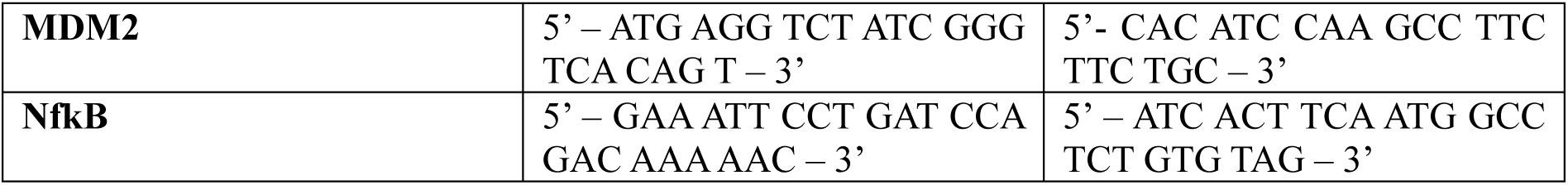

### Metabolomics – Mass spectrometry (MS) method

Healthy PCLS were incubated at 37 °C and 5% CO_2_ in 24-well plates. On certain time points (at 6h, 8h, day 1, day 2, day 3, day 4, day 7, day 10, day 11, day 14, and day 17) 10µL of supernatant was collected and stored in -80°C for further analysis. The culture media was refreshed on day 4, and day 10. For analysis, 10 µl of the sample was loaded into a Dionex UltiMate 3000 LC System (Thermo Scientific Bremen, Germany) equipped with a C-18 column (Acquity UPLC -HSS T3 1. 8 µm; 2.1 x 150 mm, Waters) coupled to a Q Exactive Orbitrap mass spectrometer (Thermo Scientific) operating in negative ion mode. A step gradient was carried out using solvent A (10 mM TBA and 15 mM acetic acid) and solvent B (100% methanol). The gradient started with 5% of solvent B and 95% of solvent A and remained at 5% B until 2 min post injection. A linear gradient to 37% B was carried out until 7 min and increased to 41% until 14 min. Between 14 and 26 minutes the gradient increased to 95% of B and remained at 95% B for 4 minutes. At 30 min the gradient returned to 5% B. The chromatography was stopped at 40 min. The flow was kept constant at 0.25 mL/min and the column was placed at 40°C throughout the analysis. The MS operated in full scan mode (m/z range: [70.0000-1050.0000]) using a spray voltage of 4.80kV, a capillary temperature of 300°C, sheath gas at 40.0, and auxiliary gas at 10.0. The AGC target was set at 3.0E+006 using a resolution of 140000, with a maximum IT fill time of 512 ms. Data collection was performed using the Xcalibur software (Thermo Scientific). The data analyses were performed by integrating the peak areas (El-Maven – Polly - Elucidata).

### Orthotopic Tumor model

Renca cells were transduced with a viral vector (EF1a-GFP-FLuc-Puro) containing the firefly luciferase gene. In this study, 15 female BALB/c mice were used (5 weeks old) (Charles River, Beerse, Brussels) to induce orthotopic lung cancer. Mice were injected intravenously with 1 x 10^6^ RencaLuc+ cells. At the beginning, 5 mice were used to prepare cancerous PCLS to characterize the cell population and gene expression levels. For the next cohort, 15 mice were divided into 3 groups (*in vivo* treatment group (n = 5), control group (n = 5), and *ex vivo* treatment group (n = 5)) to compare the *ex vivo* treatment with *in vivo* treatment.

### *In vivo* Bioluminescence Imaging (BLI)

Tumor growth was monitored twice a week, using a non-invasive bioluminescence (BLI) optical imaging system (IVIS spectrum; PerkinElmer). For each session, mice were injected intraperitoneally (IP) with 150 mg/kg body weight d-luciferin (GoldBio, USA). The mice were then anesthetized and positioned in the IVIS Spectrum. BLI images were obtained 10 minutes post-administration of d-luciferin (medium binning, f stop = 1, excitation time = 10 sec) under general anesthesia with 2% isoflurane inhalation. Regions of interest (ROI) were indicated covering the bioluminescence signal from the tumor. Images were analyzed with LIVING Image processing software (Perkin Elmer, Waltham, MA).

### *Ex vivo* Bioluminescence Imaging (BLI)

Tumor PCLS were incubated with 200ug/mL D-luciferin. BLI images were obtained 10 minutes post-incubation with d-luciferin at room temperature (medium binning, f stop = 1, excitation time = 10 sec).

### *In vivo* treatment

For the *in vivo* treatment we choose the paclitaxel (PTX, MedChemExpress, Bio-Connect, Belgium) which is a commonly used chemotherapeutic agent. After 7 days, tumor size was measured in lung tumor-bearing mice using non-invasive BLI. 10 mice in total were divided into 2 groups: group 1 = control group (PBS, n = 10); group 2 = PTX treatment (n = 10). PTX treatment was based on intraperitoneal injection with a concentration of 20 µg/g mouse ^39^ under general anesthesia with 2% isoflurane inhalation. Intraperitoneal booster injections were given every 3 days for in total of 3 times. Control animals were injected intraperitoneally with PBS at the start of the treatment. Tumor growth was monitored twice a week. Animals were kept alive for 3 days following the third treatment booster injection and then they were euthanized.

### *Ex vivo* treatment with Paclitaxel

After 7 days of I.V. injection with Renca Luc+ cells, tumor BLI was measured in lung tumor-bearing mice using non-invasive BLI. PCLS were made from 5 mice and divided into 2 groups: group 1 = control group (culture media, n = 6 slices per mouse); group 2 = PTX treatment (n = 18 slices per mouse). PTX treatment was based on exposure of PCLS to PTX at a concentration of 20 µg/g tissue. To mimic the *in vivo* clearance, the media was changed after 4h. Booster treatments were given every 3 days for in total of 3 times. Tumor growth and tissue viability were monitored twice a week using *ex vivo* BLI and AlamarBlue HS assay.

### *Ex vivo* treatment with Ag-Citrate-5nm combined with Anti-PD1

After 14 days of I.V. injection with Renca Luc+ cells, tumor BLI was measured in lung tumor-bearing mice using non-invasive BLI. PCLS were made from 5 mice and divided into 2 groups: group 1 = control group (culture media, n = 6 slices per mouse); group 2 = Ag-Citrate-5nm (nanoComposix, USA) combined with Anti-PD1 treatment (n = 18 slices per mouse). Treatment was based on exposure of PCLS to Ag-Citrate-5nm at a concentration of 2.5 µg/g tissue and Anti-PD1 (CD279, InVivoMAb anti-mouse PD1, Biocell) at a concentration of 10 µg/g tissue. To mimic the *in vivo* clearance, the media was changed after 48h. Booster treatments were given every 3 days for in total of 3 times. Tumor growth and tissue viability were monitored at every refreshment of media using *ex vivo* BLI and AlamarBlue HS assay.

### PCLS preparation for H&E staining

To see how the morphology of the PCLS was changed during the cultivation period, eosin-hematoxylin staining was performed on sections of 10 µm thick (see supplementary for more details). Images of tissue sections were acquired with the Axioscan 7 (Zeiss, Germany) and analyzed with the open-source software FIJI.

### IHC staining

10-µm cryostat sections were prepared for immunohistochemical staining. Antigen retrieval was achieved by incubation with proteinase K for 15min. Sections were washed with 1xPBS for 5min and incubated with blocking solution (1x PBS + 10% normal goat serum (NGS, ThermoFisher scientific) + 1% FBS) for 1h. The sections were washed 2 times with 1x PBS and incubated in avidin blocking solution (0.001% avidin (Sigma-aldrich, USA) in 1xPBS) for 20 minutes. The sections were washed twice with 1xPBS and incubated with biotin blocking solution (0.001% biotin (Sigma-aldrich, USA) in 1xPBS) for 20min. After washing twice with 1xPBS, sections were incubated overnight at 4°C with primary antibody (rat – anti – mouse). Endogenous peroxidases were quenched by incubating the sections in 3% hydrogen peroxide (Acros Organics, Thermofisher Scientific, Belgium) for 20 min. The sections were then washed twice with 1xPBS. Incubation with Anti-Rat-Biotin (Biotin-SP AffiniPure Goat Anti-Rat IgG (H+L), Jackson Immunoresearch, UK) was performed for 1h at room temperature (RT). After washing twice in 1xPBS, the sections were incubated with streptavidin-HRP (HRP-Conjugated Streptavidin, ThermoFisher) for 30min at RT. After washing with 1xPBS, the sections were incubated with tyramide working solution (Alexa Fluor™ 488 Tyramide SuperBoost™ Kit, goat anti-rabbit IgG, Invitrogen, ThermoFisher scientific) by following the manufacturer instructions. Stop solution (ab210900, Abcam) was added to each section for 2min after which the sections were washed 3 times with 1x PBS. Sections were incubated overnight with primary antibody (rabbit – anti – mouse). The sections were washed twice with 1xPBS and incubated with goat-anti-rabbit-polyHRP (Invitroge, ThermoFisher) for 1h at RT. After incubation, the sections were washed once with 1x PBS and incubated for 1h with tyramide working solution (Alexa Fluor™ 555 Tyramide SuperBoost™ Kit, goat anti-rabbit IgG, Invitrogen, ThermoFisher scientific) by following the manufacturer instructions. Stop reagent was added for 2min and the sections were washed 3 times with 1xPBS. Hoechst staining was added for 10min at RT and the sections were washed 2 times with 1xPBS. Mounting was performed using fluoromount aqueous mounting medium (Sigma-aldrich, USA) and the sections were covered using cover glass (Rectangular cover glasses, VWR, Belgium). The images were acquired with Axioscan (Zeiss) (Akoya Biosciences, USA) and analyzed using open-source software QuPath.

| Primary antibody | Secondary antibody |
| --- | --- |
| Neutrophil Ly-6C (Rat-anti-mouse, Bio-Rad, USA), | Anti-Rat-Biotin |
| F4/80 (Rabbit-anti-mouse, Cell Signaling Technology, Netherlands), | Goat-anti-rabbit-polyHRP |
| F4/80 (Rat-anti-mouse, Bio-Rad, USA) | Anti-Rat-Biotin |
| Anti-CD8 (Rabbit-anti-mouse, Novus Biologicals, UK) | Goat-anti-rabbit-polyHRP |
| CD90.2 monoclonal antibody (APC conjugated, eBioscience, ThermoFisher Scientific, Belgium) | - |

Combination of different antibodies:

- Neutrophil Ly-6C (Rat-anti-mouse, Bio-Rad, USA) + F4/80 (Rabbit-anti-mouse, Cell Signaling Technology, Netherlands)
- Neutrophil Ly-6C (Rat-anti-mouse, Bio-Rad, USA) + Anti-CD8 (Rabbit-anti-mouse, Novus Biologicals, UK)
- F4/80 (Rat-anti-mouse, Bio-Rad, USA) + Anti-CD8 (Rabbit-anti-mouse, Novus Biologicals, UK)

### Cryopreservation and thawing of PCLS

Given that it is possible to generate multiple PCLS from a single organ, it is crucial to optimize cryopreservation to establish biobanks for future use. The following strategies were employed: Controlled rate freezing (n = 12), conventional freezing (n=12), and snap freezing (n=12). Cryotubes containing freezing media, such as culture media mixed with 10% DMSO or FBS mixed with 10% DMSO, were used to freeze PCLS. The conventional freezing method involved placing the cryotubes in a container with isopropanol (Mr. Frosty) and then transferring them to liquid nitrogen after they had been placed in the container for 2 hours at -80 °C. For snap freezing, the cryotubes were placed directly into liquid nitrogen. In addition, controlled rate freezing was carried out using a controlled rate freezing device (Kryo 560-16, Planer, UK), where the temperature was reduced according to the following schedule: 5min at 8°C, reduced to 4 °C with 1°C/min, reduced to 0°C with 4°C/min, kept at 0°C for 5min, reduced to -100 °C with 5°C/min and then placed in LN.

To thaw the cryotubes stored in a -196°C liquid nitrogen tank for 21 days, they were first immersed in a 37 °C water bath for 30 seconds. Subsequently, the PCLS were carefully removed from the cryotubes using a fishing loop and transferred to prewarmed culture media, where they were incubated at 37 °C and 5% CO_2_ in 24-well plates for one hour before undergoing viability measurement with AlamarBlue HS.

## Results

### Optimization of PCLS cultivation conditions for maximal viability

A robust, non-invasive viability assay is crucial for maintaining the original tissue architecture and microenvironment in Precision Cut Lung Slices (PCLS). Such an assay allows for longitudinal monitoring, providing insights into the PCTS’s metabolic activity, cytotoxicity, and ATP production. In our study, we evaluated the effectiveness of three different viability assays.

The ATP assay was used to measure both intracellular and extracellular ATP, thus indicating the metabolic activity of the PCTS over time. It’s important to note that an increase in extracellular ATP often correlates with increased cellular stress, possibly due to oxidative stress pathways (**figure 1A**). Alongside this, the Lactate Dehydrogenase (LDH) assay was employed as a cytotoxicity measure, evaluating the membrane integrity of the cells within the PCTS. Elevated levels in this assay, similar to increased extracellular ATP, typically point towards enhanced cytotoxicity(**figure 1A,B**).

**Figure 1:**
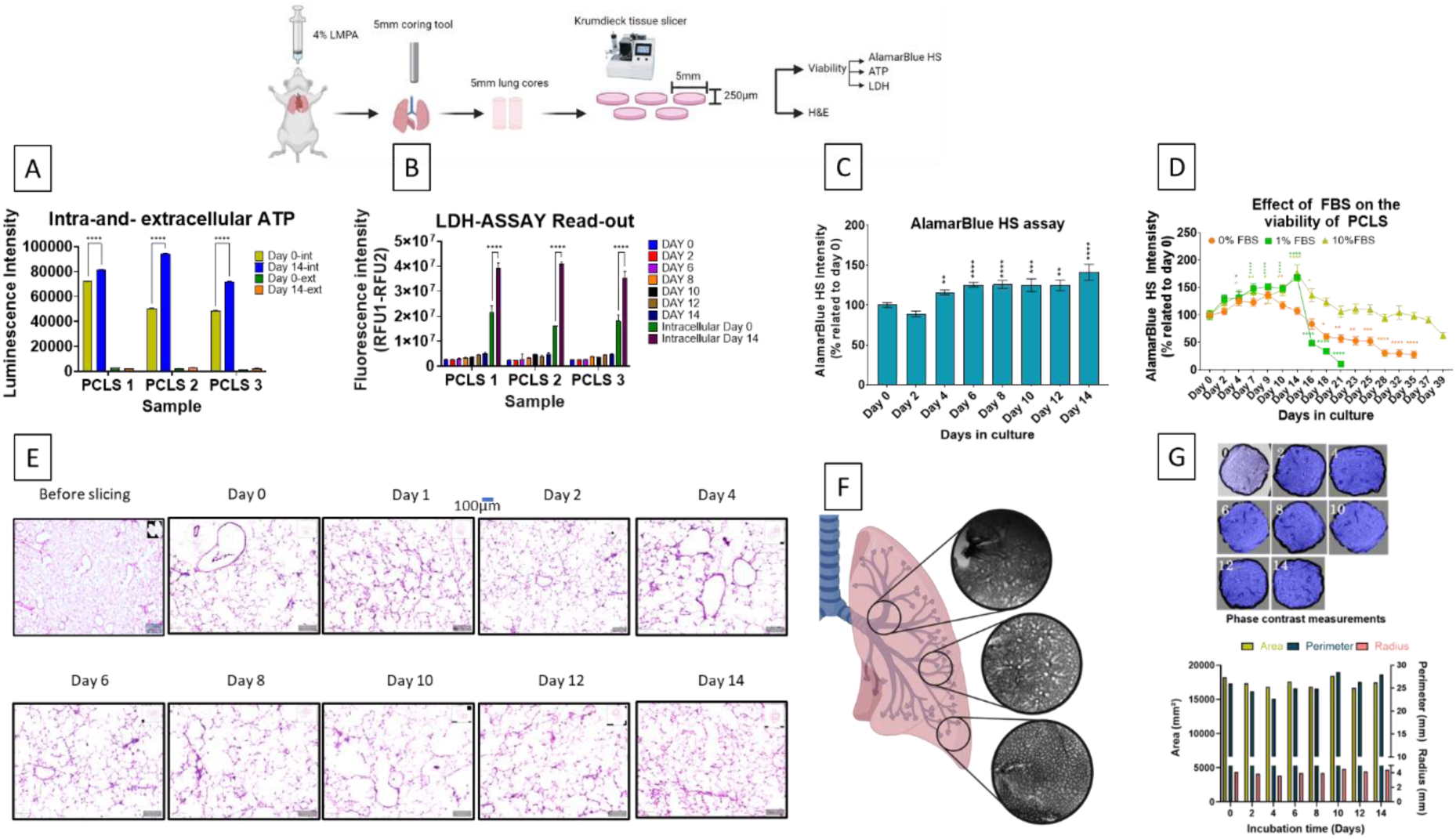
Evaluation of Viability and Structural Integrity in Precision Cut Lung Slices (PCLS) A) Intracellular and extracellular ATP measurements showing metabolic activity and cellular stress over time. B) LDH assay results indicating cytotoxicity levels within the PCLS. C) ABHS assay depicting metabolic activity recovery over 14 days post-slicing. D) Impact of varying FBS concentrations (0%, 1%, 10%) on PCLS metabolic activity, highlighting the optimal condition with 10% FBS. E) H&E staining confirming the preservation of lung structures. F-G) Phase contrast images showing consistent PCLS architecture (generated using Biorender) and size throughout the 14-day incubation period, indicating no tissue disintegration. The level of significance was indicated when appropriate (*:p< 0.05; **:p< 0.01; ***:p < 0.001; ****:p<0.0001).

Furthermore, the Alamar Blue HS (ABHS) assay was utilized to indicate the metabolic state of the tissue by measuring the reduction capacity in PCLS over time (**figure 1C**). The use of these assays in combination was instrumental; for instance, a combination of high extracellular ATP and LDH values usually indicates cytotoxicity, which is often paralleled by low ABHS reduction values in the PCLS.

After assessing each assay for ease of use, sensitivity, and invasiveness on PCLS, ABHS was chosen for further experiments. Its selection was based on its user-friendliness and non-invasive nature, allowing for the continuous monitoring of PCLS metabolism over an extended period without inducing toxicity.

The results from the ABHS assay showed that post-slicing the metabolic activity of the PCLS was low but restored after 4 days, as depicted in **(figure 1C).** This recovery suggests that the PCLS requires a period to adapt to the new culturing media and recover from the damage that occurred during slicing. Once this adaptation phase is completed, the PCLS remained metabolically active over 14 days (**Figure 1C**), demonstrating their sustained viability under our optimized culturing conditions.

Next, we wanted to investigate the impact of FBS concentration on the metabolic activity of PCLS during cultivation (**figure 1D**). To do this, we utilized culture media containing FBS at concentrations of 0%, 1%, and 10% to cultivate the PCLS. Our findings revealed that 10% FBS was effective in maintaining the PCLS’s metabolic activity for an extended period of time. Therefore, we decided to use culture media with 10% FBS for subsequent experiments.

### Long-term culture of PCLS maintains structural integrity

The preservation of architectural integrity in the PCLS throughout this duration was consistently confirmed through non-invasive PCLS imaging via phase contrast imaging at various time points as depicted in **figure 1F**. Notably, there were no signs of tissue size reduction or disintegration observed throughout the incubation period. The downstream processing of the phase contrast images confirmed that the area, perimeter, and radius of the PCLS remained consistent over 14 days of incubation (**Supplementary Figure**). The PCLS preserved the distinct lung structures such as the bronchus, bronchioles, terminal bronchiole, alveolar sac, and alveoli, as indicated in **figure 1F** Depending on the position of tissue cores these different lung structures are present on the PCLS (**figure 1F**). Furthermore, when tissue cores are taken from the upper section of the lungs, the pulmonary arteries and veins are visible on the phase contrast images while in the mid and lower section structures such as terminal bronchiole, alveolar sac, and alveoli are visible **(figure 1F).** These findings were further confirmed by H&E staining (**figure 1E**), which also affirmed the maintenance of tissue integrity and the presence of the above mentioned different lung structures. The staining vividly highlighted the distinct morphological features of each structure, ensuring that not only were these components preserved, but they also retained their functional characteristics, essential for accurate *ex-vivo* studies.

### Metabolomics improves optimized culture conditions of PCLS

To further understand the low metabolic activity at the beginning and the recovery after 4 days, we looked at the change of metabolites during the cultivation time of the PCLSs, therefore LC-MS/MS was performed on the supernatant (tissue suspension). As depicted in **figure 2A** the results demonstrated an elevated production of L-lactic acid, L-alanine, and L-proline over time relative to the control of the day of sampling. Furthermore, there was a reduced L-arginine production relative to the control of day of sampling **(figure 2A-E).**

**Figure 2:**
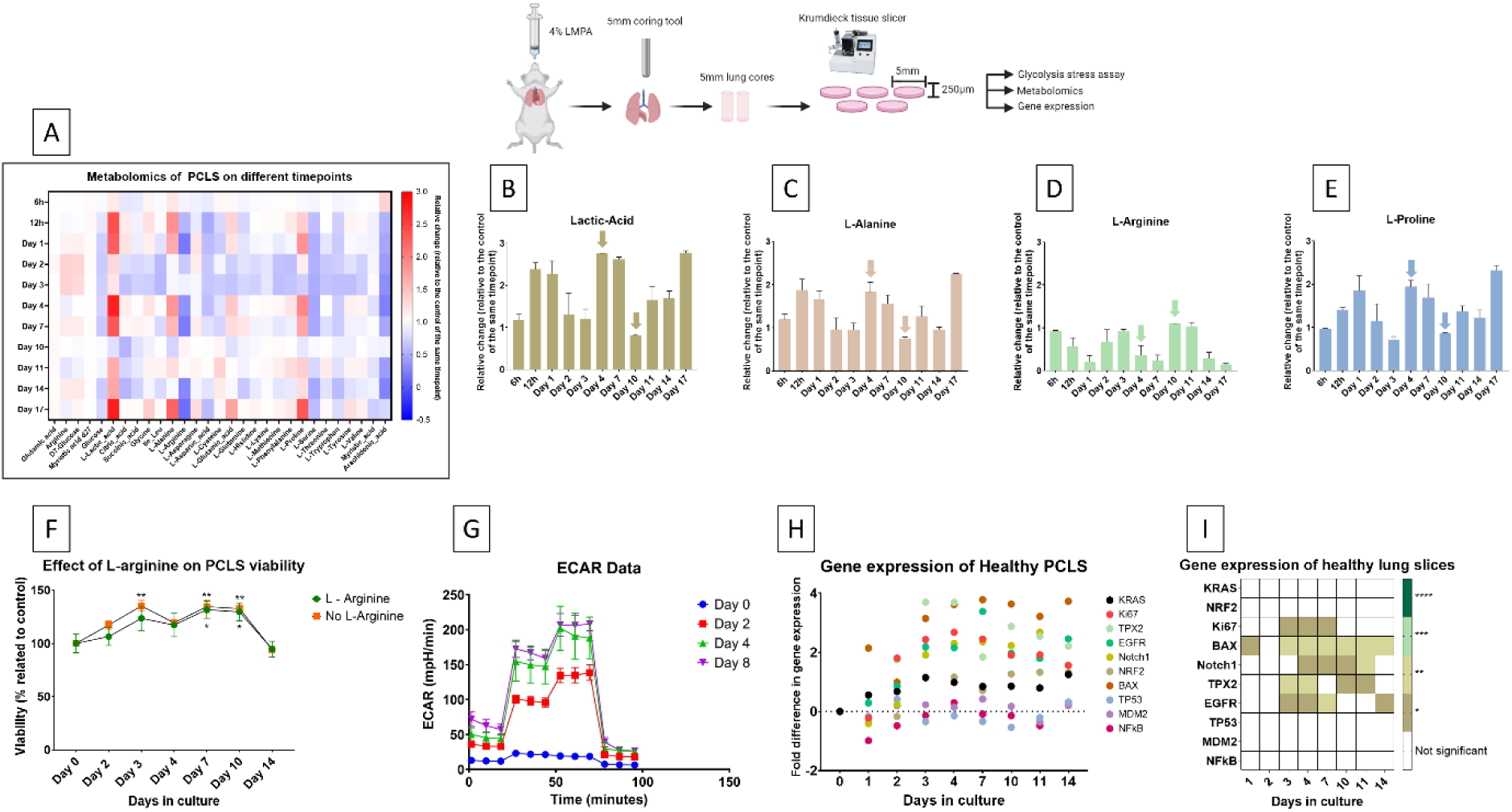
Metabolic Activity, Glycolysis, and Gene Expression in Precision Cut Lung Slices (PCLS). A-E) LC-MS/MS analysis of the supernatant showing increased production of L-lactic acid, L-alanine, and L-proline, and reduced L-arginine over time. The arrows indicate the timepoints when the media is been refreshed. F) Viability of PCLS with and without L-arginine supplementation, showing no significant improvement in the initial two days but a significant difference from day 3 onward. G) Glycolysis assay measuring ECAR, indicating increased glycolytic activity and tissue adaptation over time. H-I) Gene expression analysis showing significant changes in Ki67, BAX, Notch1, TPX2, and EGFR, with no notable alterations in KRAS, NRF2, TP53, MDM2, and NF-κB, highlighting dynamic responses in proliferation, apoptosis, differentiation, and growth signaling pathways. The level of significance was indicated when appropriate (*:p< 0.05; **:p< 0.01; ***:p < 0.001; ****:p<0.0001).

Following these results, to elucidate the effect of L-arginine on the viability of precision-cut lung slices (PCLS), the slices were incubated in media either supplemented with or without L-arginine. Comparative analysis of the results from the initial two days of cultivation indicated that L-arginine supplementation did not significantly improve the viability of PCLS when compared with those cultured in its absence (**figure 2F**). Viability assessments conducted on days 3, 7, and 10 revealed a statistically significant difference (**p-value < 0.05**) in the viability of PCLS with or without L-arginine supplementation.

To elucidate the mechanisms underlying the elevated production of L-lactic acid within the PCLSs, we conducted a glycolysis assay, measuring the extracellular acidification rate (ECAR). Our investigation revealed that immediately after slicing, the tissue exhibited the lowest ECAR levels, indicating the necessity for tissue recovery and adaptation to the new media (**figure 2G**). Initially, we observed diminished glycolytic activity at baseline (day 0), which gradually increased over the incubation period of PCLSs and reached its peak on day 8. Furthermore, our results indicate that after two days of incubation, the tissue began to manifest elevated glycolytic activity, followed by a stable glycolytic state post-day 4, signifying the attainment of tissue homeostasis.

The observed increase in glycolytic activity, as indicated by the rise in ECAR levels, correlates well with the metabolomics findings, which showed elevated production of L-lactic acid over time. This suggests that the heightened glycolytic activity contributes to the increased synthesis of this metabolite within the PCLSs. Under stress or hypoxic conditions, the PCLSs shift to highly glycolytic metabolism, where pyruvate is preferentially converted to L-lactic acid to generate energy, rather than undergoing further metabolism in the Krebs cycle and electron transport chain.

### Gene expression indicates minor adaptations to proliferation and cell survival

#### Gene Expression in Healthy PCLS

The results of the gene expression assay using precision-cut lung slices showed varying patterns of expression at different time points throughout the culture period. The expression of the Ki67 gene, a marker for cell proliferation, was significantly increased on days 3, 4, 7, and 10, indicating periods of active cell division ^40^. On the other hand, the expression of the BAX gene, which is associated with apoptosis, was significantly elevated on days 3, 4, 7, 10, 11, and 14, suggesting increased apoptotic activity necessary for removing damaged cells and maintaining tissue balance ^41^ (**figure 2H,I).**

Notch1, a critical regulator of cell differentiation and tissue remodeling, displayed elevated expression levels on days 4, 7, and 11, indicating its involvement in directing cell fate decisions during lung tissue regeneration ^42^. The **TPX2** gene, which is essential for spindle formation during mitosis, exhibited sustained elevated expression levels on days 2, 3, 4, 7, 10, 11, and 14, suggesting ongoing mitotic activity that is crucial for continuous cell division and effective tissue repair ^43^. Similarly, **EGFR**, which promotes cell growth, survival, and proliferation, was significantly upregulated on days 3, 4, 7, 10, and 14, emphasizing its role in providing robust proliferative and survival signals during lung tissue repair ^44^ (**figure 2H,I).**

Conversely, no notable alterations were observed in the expression levels of KRAS, NRF2, TP53, MDM2, and NF-κB. The consistent expression of KRAS suggests that the culture conditions did not activate oncogenic pathways or involve KRAS in immediate stress and repair responses ^45,46^. The lack of significant changes in NRF2 implies that the culture conditions did not induce substantial oxidative stress, or existing antioxidant defenses were sufficient ^47,48^. Similarly, the stability of TP53 expression indicates that the culture conditions did not elevate stress levels enough to activate p53-mediated pathways, with cellular repair mechanisms managing damage effectively ^49,50^. The unaltered expression of MDM2, which regulates TP53 activity, suggests a balanced regulatory environment (Wade et al., 2013). Lastly, the stable expression of NF-κB implies that the culture conditions did not provoke significant inflammatory responses or that any inflammation was transient and did not lead to sustained gene expression modifications ^51^(**figure 2H,I).**

These findings collectively highlight a dynamic and coordinated response involving proliferation, apoptosis, differentiation, and growth signaling pathways, essential for effective lung tissue repair and regeneration. The absence of significant changes in key regulatory genes indicates that the culture conditions maintained cellular homeostasis without triggering major stress or damage response pathways.

### Optimization of cryopreserved Precision-Cut Lung Slices

The purpose of this study was to assess the optimal cryopreservation method and medium by analyzing three techniques: conventional freezing, controlled-rate freezing, and snap freezing. In addition, the effectiveness of two distinct cryoprotective media formulations, comprising culture media mixed with 10% DMSO and FBS with 10% DMSO, was evaluated.

Following conventional freezing, PCLS demonstrated a prompt recovery, with metabolic activity returning to homeostatic levels within three days post-thawing (Figure). The maintained viability over 21 days post-thawing suggests this method’s potential for long-term experimental applications. The type of freezing media—either culture medium+10% DMSO or FBS+10% DMSO—appeared to have negligible impact on tissue viability (**figure 3A**).

**Figure 3:**
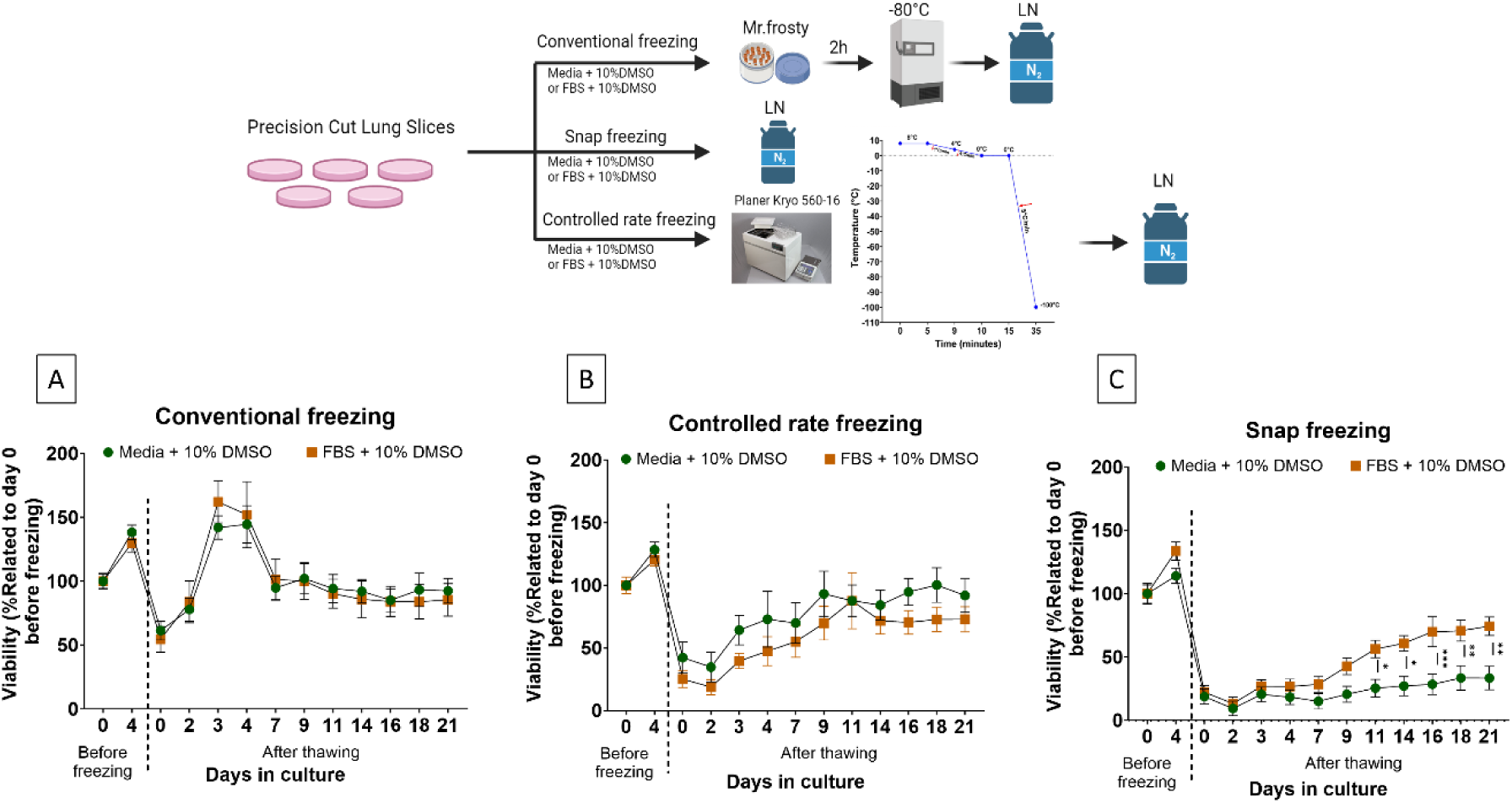
Cryopreservation Methods and Media for Precision-Cut Lung Slices (PCLS): A) Viability recovery of PCLS post-conventional freezing using culture medium + 10% DMSO and FBS + 10% DMSO, showing prompt recovery within three days and maintained viability over 21 days. B) Recovery pattern of PCLS post-controlled-rate freezing, reaching equilibrium approximately 14 days after thawing with negligible impact from the freezing media composition. C) Viability recovery of PCLS post-snap freezing, with FBS + 10% DMSO reaching 50% viability in 9 days, while culture medium + 10% DMSO failed to achieve 50% viability within the 21-day period, highlighting the significant influence of cryoprotectant choice on tissue recovery. The level of significance was indicated when appropriate (*:p< 0.05; **:p< 0.01; ***:p < 0.001; ****:p<0.0001).

Controlled-rate freezing showed a pattern of recovery similar to that of conventional freezing, with PCLS regaining equilibrium approximately 14 days after thawing (**figure 3B**). The rate of recovery was more gradual, and, as with conventional freezing, the freezing media composition did not significantly affect the post-thaw viability of the tissue slices.

However, PCLS subjected to snap freezing displayed significantly reduced recovery. It took around 9 days for the tissue frozen in FBS+10% DMSO to reach 50% viability, and the PCLS frozen in culture medium+10% DMSO did not achieve 50% viability at any point during the 21-day culture period **p < 0.01** (**figure 3C**). Notably, the choice of cryoprotectant markedly influenced tissue viability, with FBS+10% DMSO showing superior outcomes to culture medium +10% DMSO.

### Generation of cancerous PCLS and optimization of analysis methods

Balb/c mice were injected with genetically engineered Renca luciferase positive cells through the tail vein, which enabled the establishment of an *in vivo* artificial lung metastasis model. The expression of luciferase facilitated non-invasive monitoring of tumor growth through bioluminescence, where the enzyme reacts with D-Luciferin to emit light. Upon detection of tumor growth (as depicted in Figure mice in IVIS), PCLS were prepared from the mice. Upon exposure of PCLS to D-Luciferin, the presence of tumor cells could be visualized through the bioluminescence observed in a 24-well plate image (as shown in Figure 24 well plate IVIS). Moreover, H&E staining was used to validate the presence of tumor cells within the PCLS, as indicated by the cancerous region marked with circles (**figure 4A**).

**Figure 4:**
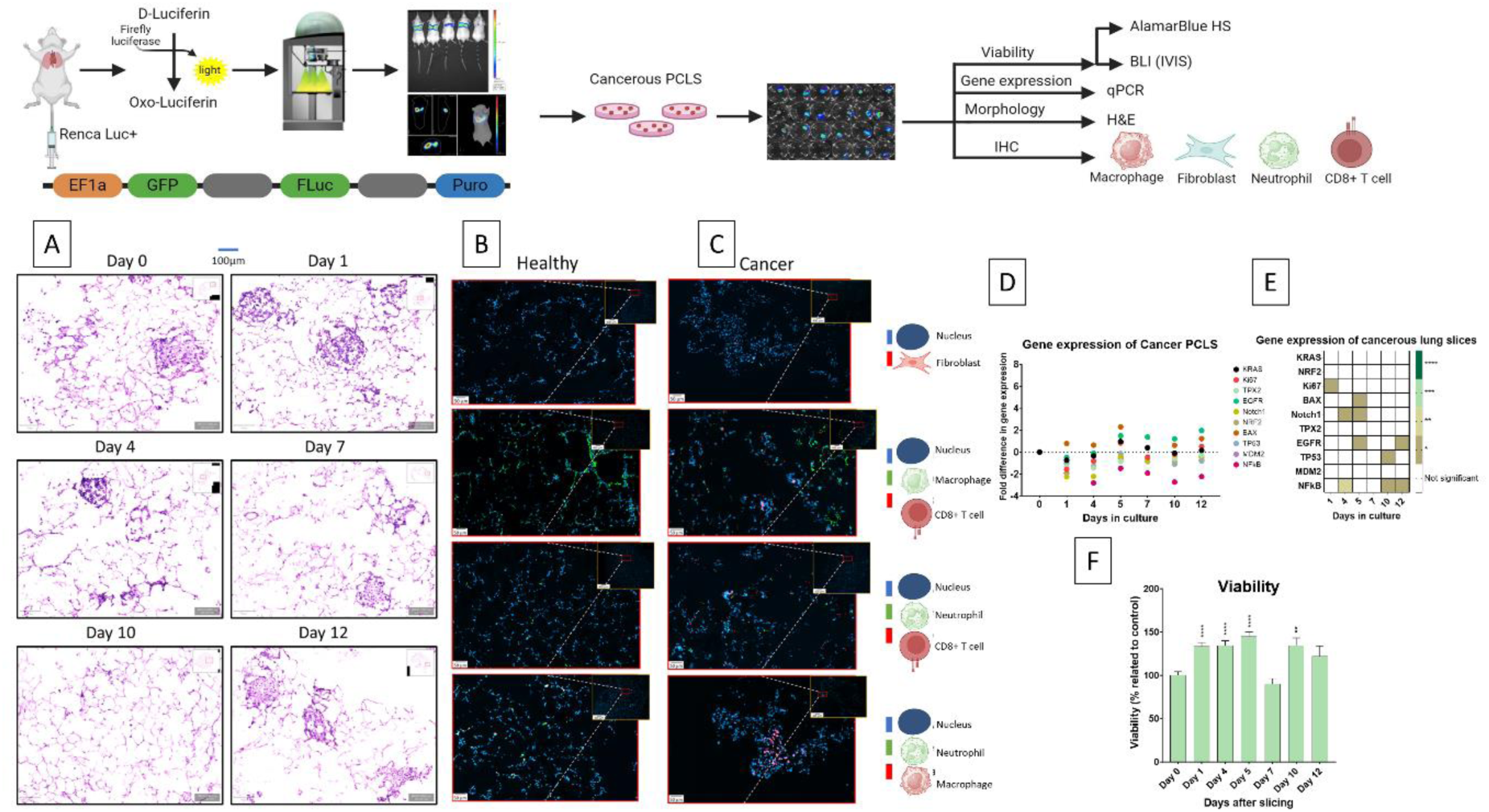
Characterization of Cancerous PCLS and Gene Expression Analysis: A) H&E staining of cancerous PCLS. B) IHC of healthy PCLS showing uniform cell nuclei distribution and minimal immune cell presence. C) IHC of cancerous PCLS indicating increased nuclear density and significant infiltration of fibroblasts, macrophages, neutrophils, and CD8+ T cells. D-E) Gene expression analysis in cancerous PCLS highlighting elevated levels of Ki67, TPX2, BAX, Notch1, NRF2, EGFR, NF-κB, TP53, and MDM2 at various time points, with no significant change in KRAS expression. F) AlamarBlue HS assay results showing high metabolic activity in cancerous PCLS over 12 days, with significant increases on days 1, 4, 5, and 10. The level of significance was indicated when appropriate (*:p< 0.05; **:p< 0.01; ***:p < 0.001; ****:p<0.0001).

### Optimized tumor-derived PCLS retain their tumor and normal host tissue microenvironment

In the healthy PCLS, DAPI staining reveals a uniform distribution of cell nuclei with a consistent density that reflects the structured architecture of normal lung tissue (**figure 3B**). The absence or low visibility of specific immune cell markers, such as those for fibroblasts, neutrophils, and CD8+ T cells (**figure 4B)**, suggests minimal immune activity, which is characteristic of non-pathological states. There is no significant indication of inflammation or fibroblast activation, corroborating the expected cellular composition and microenvironment of healthy lung tissue. The IHC showed the presence of residual macrophages.

In contrast, the cancerous PCLS exhibits a notably different staining pattern. DAPI staining shows an increased density of clustered nuclei, indicating a disrupted architecture and likely an enhanced proliferation rate. Notably, there is a prominent presence of fibroblasts, which are often activated in response to tumor growth (**figure 4C)**. The marked infiltration of macrophages, neutrophils, and CD8+ T cells (**Figure 4C)**, as highlighted by their respective markers, points to a robust immune response, a hallmark of the tumor microenvironment. This immune infiltration is indicative of the dynamic interactions between the tumor and the host’s immune system within the cancerous PCLS.

These findings demonstrate that PCLS effectively preserve lung architecture as well as the distinct cell populations of healthy and cancerous tissues, validating the model for studying both physiological and pathological states.

### Viability and gene expression of cancerous PCLS demonstrate optimal culture and minor adaptations to cell survival and proliferation

AlamarBlue HS assay was performed on cancerous PCLS. The metabolic activity stayed high during the cultivation period of 12 days. Significant increase in metabolic activity was observed at day 1, 4, 5, and 10 (**figure 4F)**.

### **Gene Expression of Cancer PCLS (****figure** 4D,E)

Our research investigated the gene expression patterns in PCLS from mice that had been injected intravenously with renal adenocarcinoma (Renca) cells 14 days prior to the preparation of the PCLS. The gene expression profiles showed substantial changes in several genes that are known to be involved in tumor progression and metastasis over time. **Ki67** is a marker of cellular proliferation that showed a significantly higher expression on days 1, 4, and 7 post-slicing, indicating active proliferation of metastatic cells during these early stages ^40^. **TPX2**, which is essential for spindle assembly during mitosis^52^, was significantly elevated on days 1 and 4, suggesting robust cell division activity. The pro-apoptotic gene **BAX** showed increased expression across multiple time points (days 1, 4, 5, 7, 10, and 12), indicating the induction of apoptotic mechanisms ^41^.**Notch1**, which plays a crucial role in cell differentiation and fate determination, was found to be significantly upregulated on days 1 and 4. This suggests that it has an early and significant role in cellular processes ^53^. Similarly, **NRF2**, which is responsible for regulating the antioxidant response, exhibited a significant upregulation on days 1 and 4, suggesting a response to oxidative stress ^47^. **EGFR**, a crucial receptor tyrosine kinase that promotes cell survival and proliferation, displayed notably heightened expression on days 5, 7, 10, and 12, implying its role in sustaining long-term metastatic growth ^44^. **NF-κB**, a central mediator of inflammation and cell survival, exhibited persistent activation on days 1, 4, 5, 7, 10, and 12, which suggests its role in fostering a pro-inflammatory and pro-survival microenvironment ^54^. The expression of the tumor suppressor gene **TP53**, which plays a crucial role in DNA repair and apoptosis, was found to be higher on days 4 and 10, indicating its regulatory function during these specific periods ^49^. **MDM2**, a negative regulator of TP53, displayed a significant upregulation on day 4, thereby indicating its involvement in modulating TP53 activity ^50^. No noticeable alterations in the expression of **KRAS** were detected across the various time points investigated.

### PCLS as a comprehensive model for *in vivo* experiments

In this study, we aimed to assess the representativity of PCLS for modeling metastatic lung cancer, using a well-established Renca Luc+ Balb/c mouse model. By administration of the Renca cells into the tail vein, an artificial pulmonary metastasis model is created. Upon confirmation of tumor presence in the lungs, mice were divided into two main groups for comparative analysis: one subjected to *in vivo* analysis and the other to *ex vivo* evaluation using PCLS. Each group was further subdivided into control (PBS for *in vivo* and media change for *ex vivo*) and treatment arms, with the latter receiving paclitaxel (PTX). The dose of PTX for the PCLS was kept the same as the *in vivo* dosage (20µg/g tissue), allowing for a direct comparison of treatment effects between the two models.

Subsequent to the administration of Renca Luc^+^ cells, clearly detectable lung tumor growth was identified via bioluminescence imaging seven days post-injection, as illustrated in **figure 5A**. At this critical point, mice were divided into their respective groups. The treatment protocol was initiated seven days after Renca cell injection, with the treatment group receiving paclitaxel (PTX) at 20µg/g mouse body weight. The regimen included subsequent boost doses on days 11 and 14. The control group was administered identical volume of phosphate-buffered saline (PBS), following an identical schedule to ensure methodological consistency. This strategic initiation and continuation of treatment were pivotal for evaluating the comparative efficacy of PTX against tumor progression.

**Figure 5:**
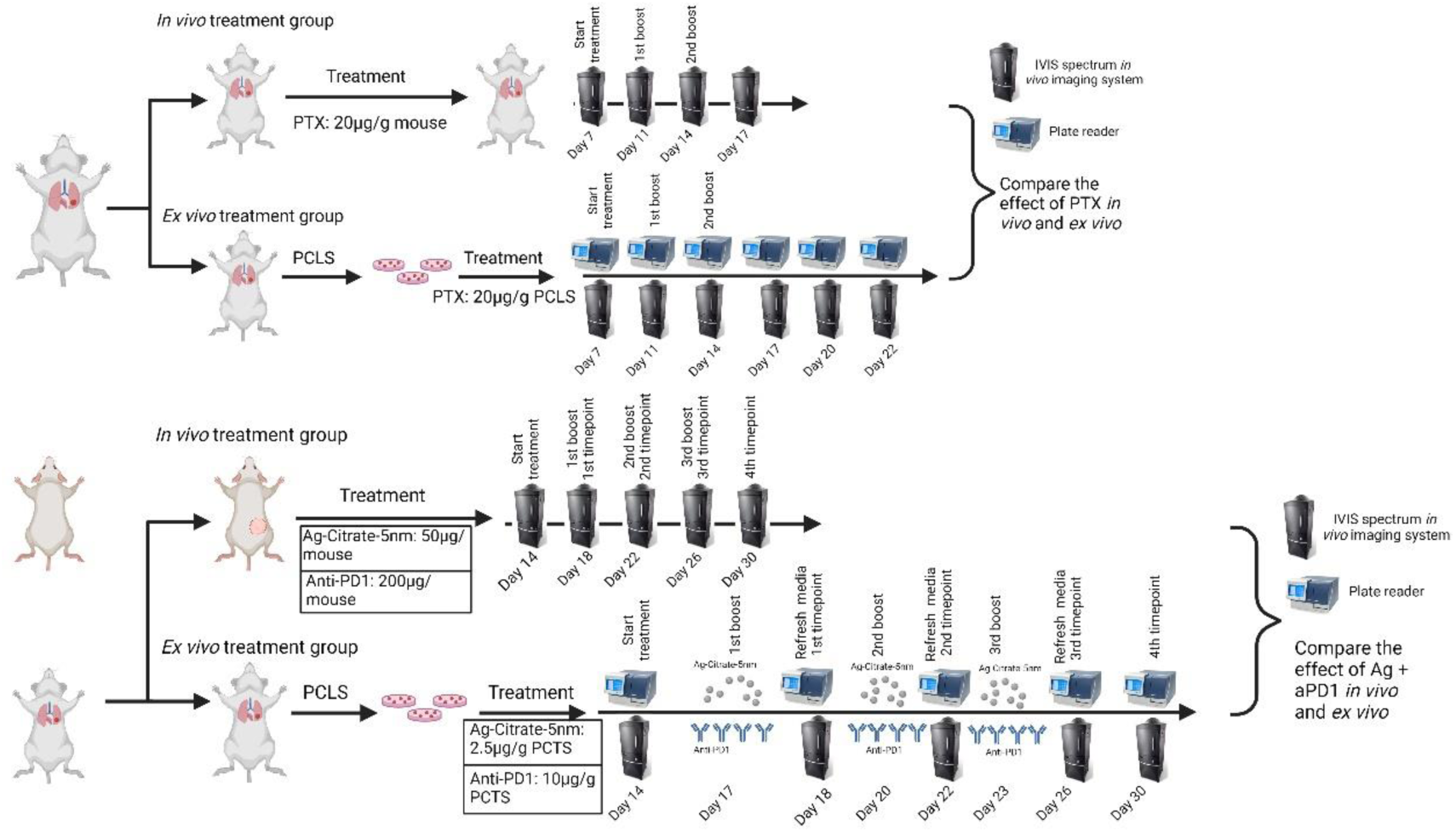
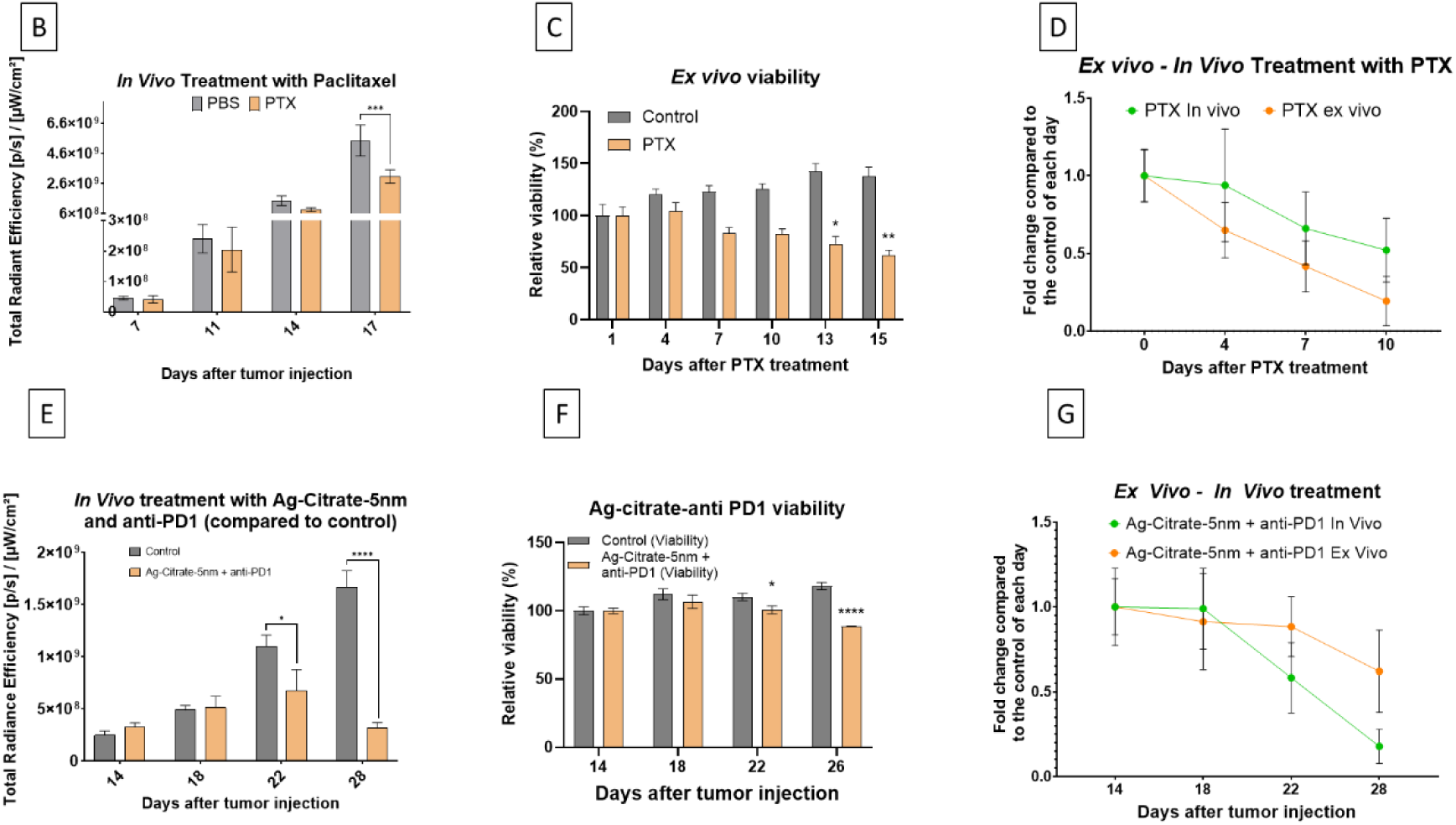
Comparative analysis of Paclitaxel (PTX) and Ag citrate NP treatment in *in vivo* and ex vivo models using precision-cut lung slices (PCLS): A) Artificial metastasis was induced in mice using Renca Luc+ cells and divided into two groups: *in vivo* treatment with Paclitaxel, monitored by IVIS, and *ex vivo* treatment, monitored by IVIS and plate reader for viability readings. Ag-Citrate-5nm treated group in vivo results were obtained in a previous study^55^ and *ex vivo* PCLSs were treated with Ag-Citrate-5nm and Anti-PD1. B) *In vivo* tumor size monitoring via bioluminescence illustrating tumor growth retention in the PTX-treated group compared to the PBS control group with significant differences observed by day 17 post-injection. C) *Ex vivo* analysis of tumor-bearing PCLS, comparing bioluminescence signal and tissue viability between PTX-treated and control groups. The PTX-treated PCLS showed a continuous reduction in tumor signal and viability, whereas the control group initially exhibited a transient reduction in tumor burden followed by resumed growth and late-stage tissue viability decline. D) Comparative evaluation of PTX therapeutic efficacy *in vivo* and ex vivo, demonstrating similar trends in tumor growth retention, supporting the relevance of PCLS as an ex vivo model for drug studies. E) Results of our previous findings showing tumor flux of Renca Luc+ cells in photon per flux. Subcutaneous Renca tumors were treated with Ag-citrate-5nm (50μg/mouse) and Anti-PD1 (200μg/mouse) with significant differences observed at 3^rd^ and 4^th^ timepoints. F) Tumor flux of Renca Luc+ cells in photon per flux and fluorescence intensity of AlamarBlue HS of PCTS. PCTS containing Renca Luc+ cells were treated with Ag-citrate-5nm (2,5μg/g PCTS) and Anti-PD1 (10μg/g PCTS) with significant differences observed at 3^rd^ and 4^th^ timepoints compared to control. G) Comparing the effect of Ag-Citrate-5nm and Anti-PD1 treatment *in vivo* and *ex vivo* relative to the control of the same timepoint. The level of significance was indicated when appropriate (*:p< 0.05; **:p< 0.01; ***:p < 0.001; ****:p<0.0001).

Simultaneously, for *ex vivo* analysis, mice were euthanized seven days post-injection, and precision-cut lung slices (PCLS) were prepared from the tumors. The PCLS were then subjected to a comparable PTX treatment regimen, receiving doses equivalent to the *in vivo* weight-adjusted dosages, to directly compare the treatment effects across both models. Alongside, a control group for the PCLS was established, where the media was refreshed at each treatment point, serving as a baseline to evaluate the effect of PTX against the untreated control condition.

Following the initial treatment phase, observations between days 7 and 14 post-injection did not reveal a significant change in tumor size between the control and treatment groups *in vivo* (**figure 5B**). Tumor growth was monitored using bioluminescence, and expressed as total radiant efficiency ([p/s]/[µW/cm^2^]). Although there was no significant difference in tumor growth rates during this period, the data suggested a trend towards tumor growth retention in the PTX-treated group compared to controls. As the study progressed, by day 17 post-injection, a significant difference was observed between the PTX treatment group and the control group *in vivo* (**figure 5B**). The PTX-treated mice exhibited retained tumor growth, whereas the PBS-treated control group showed an increased rate of tumor growth. This indicates the efficacy of PTX treatment in halting tumor progression compared to the untreated condition.

The tumor-bearing PCLS were categorized into two groups for the investigation: a treatment group and a control group. The treatment group received PTX, while the control group underwent media changes without drug administration, serving as a baseline for comparison. To assess tumor size within these PCLS, total radiant efficiency of bioluminescence was measured, providing a quantitative marker of tumor burden.

In addition to tumor growth, the overall tissue viability was meticulously monitored over time using the Alamar Blue assay. This step was critical to ensure the experimental validity, as evaluating the efficacy of the treatment on non-viable tissue would yield irrelevant results. The viability assessment allowed for the confirmation that observed responses to treatment were reflective of active biological processes within live tissue, ensuring the relevance and accuracy of the therapeutic efficacy evaluation.

In contrast, the control group exhibited an initial decrease in bioluminescence signal within the first five days, suggesting a transient reduction in tumor burden. However, this was followed by a gradual increase in signal intensity, indicative of tumor growth resumption while the PCLS treated with PTX showed continue reduction in bioluminescence signal (**figure 5C**). This delineates a clear divergence in tumor progression dynamics between the treated and control cohorts.

Concurrent viability assays, aimed at evaluating the overall health of the tissue, paralleled the trends observed in tumor signal measurements. PCLS from the treatment group displayed a decrease in viability, which was in line with the observed reduction in tumor burden (**figure 5C**). This trend suggests that the tumor reduction was associated with a corresponding decline in viable tumor cells. In contrast, the control group’s tissue viability showed a slight upward trend, culminating in a plateau that was maintained during the incubation period. However, a notable drop in viability was recorded on day 22, the final day of the experiment (**figure 5C**), signaling a late-stage decline in tissue health.

Subsequently, the therapeutic efficacy of PTX was evaluated both *in vivo* and ex vivo to determine the relevance of PCLS as an ex vivo model for studying drug effects, with the aim of reducing animal usage and minimizing experimental variation arising from inter-animal differences. The side-by-side analysis of PTX treatment outcomes revealed that PCLS can serve as a suitable ex vivo surrogate for *in vivo* conditions (**figure 5D**). Although the tumor growth retention curves for *in vivo* and ex vivo studies were not identical, they exhibited comparable trends, and the rates of decline were similar **(figure 5D).**

According to our previous research findings published in the Journal of Nanobiotechnology ^55^, the *in vivo* treatment of subcutaneous Renca Luc+ tumors with Ag-Citrate-5nm in combination with Anti-PD1 significantly decreased tumor growth rates at the third and fourth time points as compared to controls (**figure 5E)**. Similarly, the *ex vivo* treatment of PCTS derived from artificially induced lung tumors using Renca Luc+ cells also resulted in a significant reduction in tumor growth rates at the third and fourth time points (**figure 5F**, p < 0.05). The fold change analysis exhibited a consistent pattern of tumor reduction in both *in vivo* and *ex vivo* treatments (**figure 5G)**. Furthermore, the viability of PCTS remained stable, indicating that the antitumor effects were not due to general cytotoxicity (**figure 5F)**.

## Discussion

In our study, we aimed to evaluate the effectiveness of various viability assays for assessing the viability and metabolic activity of PCLS. PCLS represent a valuable *ex vivo* model system for studying lung physiology and pathology, necessitating the development of robust and non-invasive viability assays to maintain tissue integrity and functionality over time.

The combination of ATP, LDH, and ABHS assays provided comprehensive insights into PCLS viability, metabolic activity, and cytotoxicity. The ATP assay, measuring both intracellular and extracellular ATP levels, served as an indicator of metabolic activity. We observed that an increase in extracellular ATP often correlated with heightened cellular stress, potentially indicating oxidative stress pathways. This finding underscores the importance of monitoring ATP levels as a marker of cellular health in PCLS experiments. The limitation of this assay is that we change the supernatant every 2 days with fresh supernatant. This makes that the detection of extracellular ATP is not accurate. Also, we need to lyse the PCLS in order to measure the intracellular ATP which makes that we need different PCLS every time. Since the PCLS are from different parts of the lungs and from different depth, it makes that they might contain slightly different cell population and quantity. This makes the intracellular ATP measurement less accurate. Another disadvantage of the ATP assay is that by every measurement we lose PCLS.

Concurrently, the LDH assay proved valuable for assessing cytotoxicity by evaluating membrane integrity. Elevated LDH levels typically indicate compromised cell membrane integrity and enhanced cytotoxicity. The correlation between increased extracellular ATP and elevated LDH levels further highlights the interconnectedness of cellular stress and cytotoxicity in PCLS. The same limitations apply for the LDH assay.

Additionally, the ABHS assay provided insights into tissue metabolic state by measuring reduction capability over time. The use of these assays in combination allowed for a comprehensive assessment of PCLS viability and metabolic activity. Based on ease of use, sensitivity, and invasiveness, ABHS was selected for further experiments due to its non-invasive nature and suitability for continuous monitoring of PCLS metabolism without inducing additional stress or toxicity.

The results from the ABHS assay indicated that post-slicing, the metabolic activity of PCLS was initially low but restored after 4 days, suggesting an adaptation phase to the new culturing media. Our results are in line with the study conducted by Preuß et al. where they showed that the metabolic activity was decreased within the first 24h after PCLS generation ^56^. Subsequently, PCLS remained metabolically active over 14 days, demonstrating sustained viability under optimized culturing conditions.

To further validate these findings, additional cytotoxicity assays were performed through the measurement of extracellular LDH levels every 2 days. The results showed no significant increase in extracellular LDH levels over time, indicating preserved membrane integrity. Further analysis comparing intracellular LDH and ATP levels at day 0 and day 14 confirmed the viability of the tissue over time. The ratio of intra-to-extracellular LDH levels provided additional evidence of tissue viability and membrane integrity maintenance over the cultivation period. In order to assess the viability of the PCLS for longer time of period, we decided to continue with the ABHS assay since we always can use the same PCLS to monitor its viability overtime which gives us more accurate results.

Next, we wanted to see if the FBS concentration had an effect on the viability of the PCLS and if it can improve the long-term cultivation of the slices. We used F12/DMEM media with different concentration of FBS. The results showed that the highest viability was obtained with 10% FBS. At day 14 the viability reached its highest value, which is in line with our previous viability results. At day 39 the reduction capability of the PCLS in 10% FBS were reduced to 50% compared to day 0. PCLS in the lower percentage of FBS could be kept viable for up to 16 days after slicing. We did not test higher concentration of FBS because the higher the percentage of FBS the more the chance that the cells start to change their characteristics based on the small molecules present in the FBS ^57^. Our results are in line with the study of Nußbaum *et al*. where they kept PCLS viable over a period of 29 days ^58^. Our model showed that the reduction capability of the PCLS can be kept high over a period of 39 days. To better assess the viability, it would be necessary to also look at other parameters in order to verify the viability, functionality, characteristics and integrity of the PCLS after 39 days as the cells might change after long term cultivation as it is been shown that long term cultivation of PCLS decreased cytokine and chemokine production while PCLS cultured for 14 days still showed cytotoxic, inflammatory and immune response ^5960^. For this reason, we kept the PCLS in culture for 14 days for our next experiments.

Apart from the metabolic activity of the PCLS, the maintenance of architectural integrity in PCLS is crucial for accurately modeling lung physiology *ex vivo*. Therefore we employed non-invasive phase contrast imaging to assess the structural preservation of PCLS over a 14-day incubation period. The analysis of the phase contrast images, including measurements of area, perimeter, and radius data revealed no signs of tissue size reduction or disintegration, indicating the robust preservation of tissue architecture throughout the duration of the experiment.

Importantly, the distinct lung structures within the PCLS, including bronchi, bronchioles, terminal bronchioles, alveolar sacs, and alveoli, were well-preserved. Notably, the presence of pulmonary arteries and veins was visible in PCLS derived from upper lung sections, while structures such as terminal bronchioles, alveolar sacs, and alveoli were more prominent in mid and lower lung section-derived PCLS.

These findings were corroborated by histological analysis using H&E staining, which vividly highlighted the morphological features of each lung structure. This results are in line with the study conducted by Munyonho *et al*. where they confirmed the presence of normal lung alveolar architecture of PCLS after 14 days of culture ^61^. The preserved integrity and functionality of these structures are essential for accurately studying lung physiology and disease progression *ex vivo*, ensuring that PCLS maintain their physiological relevance as model systems for lung research.

To further understand the metabolomics during the cultivation of PCLS, we performed LC-MS/MS analysis on the tissue supernatant. Our results revealed dynamic changes in metabolite production over time, including elevated levels of L-lactic acid, L-alanine, and L-proline, and reduced L-arginine production relative to control of the day of sampling. In order to have a decent amount of metabolites in the supernatant, we did not change the media from day 0 to day 4. The media was changed at day 4 and the PCLS were kept in the same media till day 7. At day 7 we changed the media again and kept the PCLS in the same media till day 10 when we changed the media again and kept the PCLS in culture till day 14.

These metabolic alterations may reflect adaptive responses of PCLS to environmental and culturing conditions.

To specifically investigate the effect of L-arginine on PCLS viability, we conducted viability assessments on PCLS cultured in media supplemented with or without L-arginine. Our findings indicated that PCLS cultured in media without L-arginine supplementation exhibited same viability compared to those cultured in L-arginine-supplemented media. This suggests that the presence of L-arginine may not be essential for maintaining PCLS viability under the tested conditions. The reason that we do not see the same high viability value at day 14 compared to our previous viability assays is because we did not change the media every 2 days as we did before. By changing the media every 2 days, we supply the PCLS with fresh nutrients. Besides, we showed that there were elevated levels of L-lactic acid in the supernatant. Since L-lactic acid is acidic, it might reduce the pH level of the media where the PCLS are cultured and induce additional damage to the cells.

Since L-lactic acid is produced during glycolysis, we conducted a glycolysis assay, measuring the extracellular acidification rate (ECAR) as a proxy for glycolytic activity. Our investigation revealed dynamic changes in glycolytic activity over the incubation period, with the lowest ECAR levels immediately after slicing, followed by a gradual increase and stabilization of glycolytic activity. The observed increase in glycolytic activity correlated well with the metabolomics findings of elevated L-lactic acid production over time. This suggests that the heightened glycolytic activity contributes to the increased synthesis of L-lactic acid within PCLS. These metabolic adaptations may reflect the tissue’s response to environmental stressors and the need to generate energy under hypoxic or nutrient-deprived conditions.

To better understand the effect of long-term cultivation of PCLS in culture, gene expression assay is crucial. Understanding the gene expression dynamics in Precision Cut Lung Slices (PCLS) is essential for elucidating the underlying cellular processes and molecular mechanisms governing lung physiology and disease.

The findings from the study suggest that Ki67 and BAX levels increase together on days 3, 4, 7, and 10, indicating that cell proliferation and apoptosis cooperate in the restoration and regeneration of lung tissue. This dual mechanism indicates that the tissue is attempting to eliminate damaged cells (through BAX) and promote the growth of new cells (through Ki67), thereby maintaining a critical balance for tissue health, as reported by Yang *et al*^40^. and Guinee *et al*^41^. The process of proliferation and differentiation, as indicated by the coordinated expression of Ki67 and Notch1 on days 4 and 7, is crucial for tissue regeneration. According to Xu *et al*.^42^, Notch1 plays a significant role in determining cell fate during this process and works in conjunction with Ki67 to provide proliferative signals. Both TPX2 and Ki67 serve as indicators of cell proliferation, with TPX2 pointing to sustained mitotic activity, supporting the observations by Stewart *et al*.^43^ Additionally, EGFR and Ki67 co-expression on days 3, 4, 7, and 10 underscores the tissue’s need for robust proliferative and survival signals, as highlighted by Schramm *et al*.^44^ The co-regulation of BAX and Notch1 on days 4, 7, and 11 suggests an interplay between apoptosis and differentiation crucial for tissue maintenance^42^. The overlapping expression of BAX and TPX2 on days 3, 4, 7, 10, 11, and 14 indicates a regulated equilibrium between cell division and cell death, essential for maintaining tissue balance, consistent with the findings by Tian *et al*.^52^ The simultaneous activity of apoptotic (BAX) and survival/proliferative (EGFR) pathways is crucial for effective tissue repair^41,44^. The findings on days 4, 7, and 11 regarding TPX2 and Notch1 indicate a coordinated effort in cell proliferation and differentiation, which is crucial for tissue development and regeneration^42,52^. The simultaneous expression of genes associated with differentiation and proliferation on days 4 and 7 underscores the activation of pathways that promote these processes, which play a vital role in tissue regeneration. Finally, the overlap between TPX2 and EGFR expression suggests that mitotic activity is accompanied by growth and survival signals, enabling robust tissue regeneration and maintenance.

Our research findings with respect to KRAS and its function in cellular signaling, particularly in growth and oncogenesis, are consistent with recent studies. For instance, KRAS mutations, such as G12C, are prevalent in a variety of cancers and are essential for maintaining oncogenic signaling pathways ^45,46^. These mutations often result in the continuous activation of downstream pathways, including PI3K-AKT-mTOR, which are vital for cancer cell proliferation and survival ^62^. However, our observation that KRAS remained unaltered under the given culture conditions suggests that these pathways were not activated, corroborating that the culture environment did not induce oncogenic stress. Similarly, the regulation of NRF2, which is crucial for antioxidant responses, also showed no significant changes, indicating that the culture conditions did not cause substantial oxidative stress. This observation is supported by findings that NRF2 remains stable unless there is a significant oxidative challenge ^47,48^. Our results also showed that TP53 and its regulator MDM2 remained stable, suggesting effective management of cellular stress without activating p53-mediated apoptosis. This is consistent with research indicating that p53 pathways are only activated under severe stress conditions that necessitate repair or apoptosis ^49,50^ Moreover, NF-κB, a regulator of immune response and inflammation, exhibited no notable changes, aligning with the idea that the culture conditions did not provoke significant inflammatory responses. This supports existing literature that highlights the role of NF-κB in chronic inflammation rather than transient inflammatory responses ^63^.

The results highlight the intricate cellular processes of expansion, programmed cell death, maturation, and signaling that are essential for maintaining and repairing lung tissue in a cultured environment. The specific patterns of expression for Ki67, BAX, Notch1, TPX2, and EGFR indicate coordinated regulation of these processes, which is vital for effective tissue regeneration. Furthermore, the lack of notable changes in KRAS, NRF2, TP53, MDM2, and NF-κB suggests that the lung slice culture conditions used in our study were effective in maintaining cellular homeostasis without triggering significant stress or damage response pathways.

Cryopreservation is essential for biobanking and various research applications, including drug testing and disease modeling, as it reduces the need for animal use and frequent biopsies. This study examined the recovery of PCLS after cryopreservation using three methods: conventional freezing, controlled-rate freezing, and snap freezing, and compared the effectiveness of two different cryoprotective media combinations: Culture media +10% DMSO or FBS+10% DMSO. Following conventional freezing, PCLS showed rapid recovery, with cell metabolic activity returning to normal levels within three days after thawing. The PCLS reached a steady state in terms of viability observed over 21 days post-thawing indicates the potential of this method for long-term experimental applications. Notably, the choice of freezing media—appeared to have minimal impact on tissue viability, suggesting robustness across different cryoprotective formulations. Our results are in line with the findings of Patel *et al*. where they cryopreserved human precision cut lung slices using the conventional freezing method. In their study, the slices were still viable 14 days post thawing ^64^.

Controlled-rate freezing showed a recovery pattern similar to conventional freezing, with PCLS reaching equilibrium approximately 14 days after thawing. However, the rate of recovery was more gradual compared to conventional freezing. Similar to conventional freezing, the choice of freezing media did not significantly affect post-thaw viability, further emphasizing the resilience of PCLS to different cryoprotective formulations under controlled-rate freezing conditions.

In contrast, PCLS subjected to snap freezing exhibited significantly reduced recovery. It took approximately eight days for the tissue frozen in FBS+10% DMSO to reach 50% viability, while PCLS frozen in DMEM+10% DMSO failed to achieve 50% viability throughout the 21-day culture period. Importantly, the choice of cryoprotectant markedly influenced tissue viability, with FBS+10% DMSO demonstrating superior outcomes compared to culture medium +10% DMSO. Since the tissues are not directly exposed to LN but are in cryotubes containing either culture media + 10% DMSO or FBS + 10% DMSO at 37°C, they undergo stress due to rapid change of the temperature. FBS can contribute to better cell revival during cryopreservation due to its effect as cryoprotectant, prevent intracellular ice crystal formation and act as an antioxidant for induced oxidative stress which often occurs during freezing and thawing ^65–67^.

In our next study, we employed genetically engineered Renca cells expressing click beetle green luciferase (Luc+), a puromycin resistance gene, and GFP for fluorescence-activated cell sorting (FACS) to establish an *in vivo* artificial metastatic lung cancer model in Balb/c mice. Since the first organ the cells pass through are the lungs, cells will be trapped inside the lungs and the mice will develop lung cancer. Once significant tumor growth was detected through bioluminescence intensity, the mice were euthanized, and precision-cut lung slices (PCLS) were prepared from them to monitor tumor cell presence ex vivo. The bioluminescence signal detected within a 24-well plate allowed for the evaluation of cancer cell distribution and density within the tissue slices. This approach provided a continuum from *in vivo* monitoring to ex vivo assessment, enabling the characterization of tumor growth patterns and heterogeneity. The presence of tumors within the PCLS was further substantiated by histological analysis, particularly through H&E staining performed at various time points. Cancerous regions were identified by nodes or clusters of cells, characterized by the absence of typical lung structures and the presence of densely packed cells disrupting the tissue’s normal architecture. This histological validation reinforced the accuracy of bioluminescence imaging in identifying tumor presence both *in vivo* and ex vivo. Since the cancerous PCLS showed clusters of cells on H&E stained images, we performed IHC staining to better understand the composition of the clusters. To better understand the immune cell presence at the tumor site, we first stained healthy PCLS to determine the resident immune cells in the lung slices. DAPI staining of healthy PCLS revealed a uniform distribution of cell nuclei with consistent density, indicative of the structured architecture of normal lung tissue. Furthermore, the absence or low visibility of specific immune cell markers, including those for fibroblasts, neutrophils, and CD8+ T cells suggested minimal immune activity characteristic of non-pathological states. We could observe the presence of resident macrophages in the lung slices. The presence of macrophages in the lungs plays an important role in maintaining the tissue homeostasis and filtering the inhaled air ^68^. The absence of fibroblasts in healthy lung slices indicates that nor slicing procedure nor the culturing did induce fibrosis in the lung slices. These observations indicated the preservation of expected cellular composition and microenvironment in healthy lung tissue, with no significant indication of inflammation or fibroblast activation. In contrast, examination of cancerous PCLS revealed a distinct staining pattern. DAPI staining demonstrated an increased density of clustered nuclei, which we also saw in H&E staining, suggestive of disrupted tissue architecture and likely enhanced proliferation rates. The enhanced proliferation rate was also confirmed by the gene expression results where we saw increased gene expression levels compared to healthy tissue. Additionally, there was a prominent presence of fibroblasts, often activated in response to tumor growth, indicating the presence of tumor-associated stromal cells. Moreover, the marked infiltration of macrophages, neutrophils, and CD8+ T cells in the cancerous PCLS, as evidenced by their respective IHC markers, suggested a robust immune response within the tumor microenvironment. This immune infiltration underscored the dynamic interactions between the tumor and the host’s immune system, a hallmark feature of cancerous tissues.

The viability assay also showed that the PCLS remained viable for at least 12 days. Besides the immunohistochemistry, we also explored the gene expression of cancerous PCLS. We observed a higher level of gene expression variability compared to healthy PCLS, with most genes showing an upward trend in expression over time. Ki67, a widely recognized proliferation marker, demonstrated significantly greater expression levels on days 1, 4, and 7, emphasizing the active proliferation of metastatic cells during the initial stages of lung colonization. The upregulation of TPX2, a critical component of spindle assembly during mitosis, on days 1 and 4 supports this observation, indicating heightened cell division activity. On the other hand, BAX, a pro-apoptotic gene, exhibited increased expression across multiple time points (days 1, 4, 5, 7, 10, and 12), suggesting the simultaneous activation of apoptotic mechanisms. This dual expression pattern of genes underscores the intricate interplay between proliferation and apoptosis during the establishment and growth of metastatic tumors. The comparable expression patterns of Ki67 and TPX2 in proliferating tumor cells have been documented by Wang et al. ^69^, further attesting to their roles in regulating tumor growth.

Notch1 and NRF2 were both significantly upregulated on days 1 and 4. Notch1, which is known for its role in cell differentiation and fate determination, may be crucial in early metastatic cell adaptation and colonization. NRF2, which regulates the antioxidant response, likely responds to oxidative stress associated with rapid tumor cell proliferation and metabolic activity. The early expression of these genes suggests their importance in maintaining cellular homeostasis and adaptability in a new microenvironment. This is consistent with findings by Zhang et al.^70^, who demonstrated similar activation of Notch1 and NRF2 in metastatic breast cancer, indicating their roles in cellular adaptation and stress response.

The expression of EGFR, a receptor tyrosine kinase that supports cell survival and growth, increased at later stages of the study (days 5, 7, 10, and 12). This pattern of sustained expression suggests that it plays a role in promoting long-term metastatic growth and survival, potentially contributing to resistance against cellular stress and apoptosis. Similarly, NF-κB, a key mediator of inflammation and cell survival, displayed consistent activation throughout the study (days 1, 4, 5, 7, 10, and 12). This persistent upregulation indicates its involvement in creating a pro-inflammatory and pro-survival environment, which is essential for tumor progression and immune evasion. The significance of EGFR and NF-κB in sustaining tumor growth and promoting inflammation has been well-documented by studies conducted by Gross et al. ^71^ and Ebrahim et al. ^72^, which support our findings.

The TP53 and MDM2 gene expression profiles exhibit a tightly regulated feedback mechanism. TP53, a crucial tumor suppressor involved in DNA repair and programmed cell death, displayed heightened expression on days 4 and 10. MDM2, a negative regulator of TP53, was significantly upregulated on day 4, suggesting a feedback mechanism to control TP53 activity and prevent excessive apoptosis. This regulatory interplay is critical for preserving cellular integrity and responding to DNA damage, a common characteristic of rapidly dividing tumor cells. Our results align with those of Ghafoor et al.^73^ and Levine et al.^50^, who emphasized the significance of the TP53-MDM2 axis in modulating the cell cycle and apoptosis in cancer cells. KRAS, a gene that is frequently mutated in various types of cancer and plays a role in cell signaling pathways that regulate cell growth and differentiation, did not show any significant changes in expression across the different time points that were studied. The absence of significant changes in KRAS expression may suggest that its activity is either consistently present or not critical in this specific model of metastasis.

At this point we showed that we were able to maintain the tumor microenvironment and the gene expression levels of the cancerous lung after slicing. Next, we want to see how comparable our ex vivo model is compared to *in vivo* model. In order to do this, we injected the Renca Luc+ cells I.V. in mice (n = 10) and divided the mice in 2 groups: *in vivo* treatment group (n = 5) where we will treat the mice with PTX through I.P. injection and ex vivo treatment group (n = 5) where we will expose the cancerous PCLS to PTX by adding the same dosage of PTX in media. Based on the literature study, we used the concentration of 20µg/g for PTX ^74^, for both *in vivo* and ex vivo treatment. In order to compare the effect of PTX *in vivo* and ex vivo, we took the fold change of bioluminescence of the PTX group compared to the control for the same timepoint. Although the ex vivo treatment showed greater reduction in tumor growth rate, the result showed that in both cases, we saw the same trend in decrease. The reason that ex vivo treatment showed greater reduction might be due to the fact that the cells were more exposed to PTX compared to *in vivo* situation where there is greater biodistribution and elimination of the drug even though we kept the elimination of the drug in count by changing the media with fresh media after 4h of exposure. In order to see the effect on the viability of the tissues, we also performed viability assay using ABHS. Notably, viability assays mirrored the trends observed in bioluminescence measurements, confirming that observed responses to treatment were reflective of active biological processes within live tissue. The reduction of the viability assay was not with the same rate as observed in the bioluminescence measurement due to the fact that ABHS measures the viability of all the cells present in the PCLS, including the healthy cells, while the bioluminescence is only been measure from the cells containing the luciferase gene.

Our previous *in vivo* study demonstrated a notable decrease in tumor growth rate in subcutaneous Renca Luc+ tumors following treatment with Ag-Citrate-5nm in combination with Anti-PD1. Consequently, we decided to induce artificial metastasis in the lungs using the same cell line to generate Precision Cut Lung Slices (PCLS) and treat them with Ag-Citrate-5nm at a concentration of 2.5 µg/g in combination with Anti-PD1 at a concentration of 10 µg/g. To maintain the same experimental conditions as our *in vivo* study, we allowed the tumors to grow for 14 days before creating PCLS and initiating treatment. After 48 hours of treatment, we replaced the media with fresh media to mimic *in vivo* clearance. The results of our study demonstrated that the combination treatment of Ag-Citrate-5nm and Anti-PD1 showed significant efficacy in inhibiting tumor growth in both *in vivo* and ex vivo models of Renca tumors. In our previous study published in JNB, we showed that the combination treatment of subcutaneously induced Renca tumors with Ag-Citrate-5nm and Anti-PD1 resulted in a substantial reduction in tumor growth rates at the third and fourth time points compared to the control group. Similarly, the treatment of the PCTS also showed a considerable reduction in tumor growth rates at the third and fourth time points. Moreover, the viability of PCTS remained consistent, suggesting that the observed antitumor effects were not due to general cytotoxicity but rather to the specific efficacy of the treatment combination.

The outcomes of both *in vivo* and ex vivo treatments demonstrate a consistent trend of tumor reduction, as indicated by the fold change calculations in relation to the control on the same day. This parallel reduction trend emphasizes the potential of the ex vivo PCTS model to predict *in vivo* therapeutic outcomes, showcasing the robustness and reproducibility of the treatment effect. These results corroborate the hypothesis that the combination of Ag-Citrate-5nm and Anti-PD1 can effectively reduce tumor growth through a mechanism that can be replicated in both *in vivo* and ex vivo settings. Moreover, the preservation of PCTS viability suggests that this combination therapy not only proves effective but also selectively targets tumor cells without adversely affecting the overall tissue health.

### Conclusion

In conclusion, our study highlights that PCLS can be cultured in culture media at 37°C and 5% CO2 for at least 14 days without an impact on tissue morphology. Both conventional and controlled-rate freezing methods demonstrated comparable efficacy in preserving PCLS viability post-thawing, with minimal impact from the choice of cryoprotective media. However, snap freezing resulted in substantially lower tissue viability, highlighting its limited suitability for preserving PCLS for downstream applications. These findings underscore the importance of selecting appropriate cryopreservation methods and cryoprotective formulations to maintain the integrity and functionality of PCLS, thereby facilitating their utilization in various research endeavors, including drug screening and disease modeling.

Besides, our findings also demonstrate the effectiveness of PCLS in preserving lung architecture and distinct cell populations in both healthy and cancerous tissues. The observed differences in staining patterns between healthy and cancerous PCLS validate the utility of this model for studying physiological and pathological states, offering valuable insights into lung tissue biology and disease mechanisms. Metabolic and gene expression analyses indicated adaptive responses and coordinated regulation of proliferation, apoptosis, and differentiation during cultivation.

As last, our study highlights the potential of PCLS as a representative ex vivo model for studying orthotopic lung cancer and evaluating therapeutic interventions. By providing a platform for assessing drug effects in a controlled and reproducible environment, PCLS offer a valuable alternative to traditional *in vivo* experiments, contributing to the refinement of preclinical research methodologies and the reduction of animal usage.

## ACKNOWLEDGEMENTS

This work was supported by the KU Leuven C3/20/090, C3/21/035, C24/18/101, and IDN/21/013 fundings acquired by Dr. Bella B. Manshian.

---

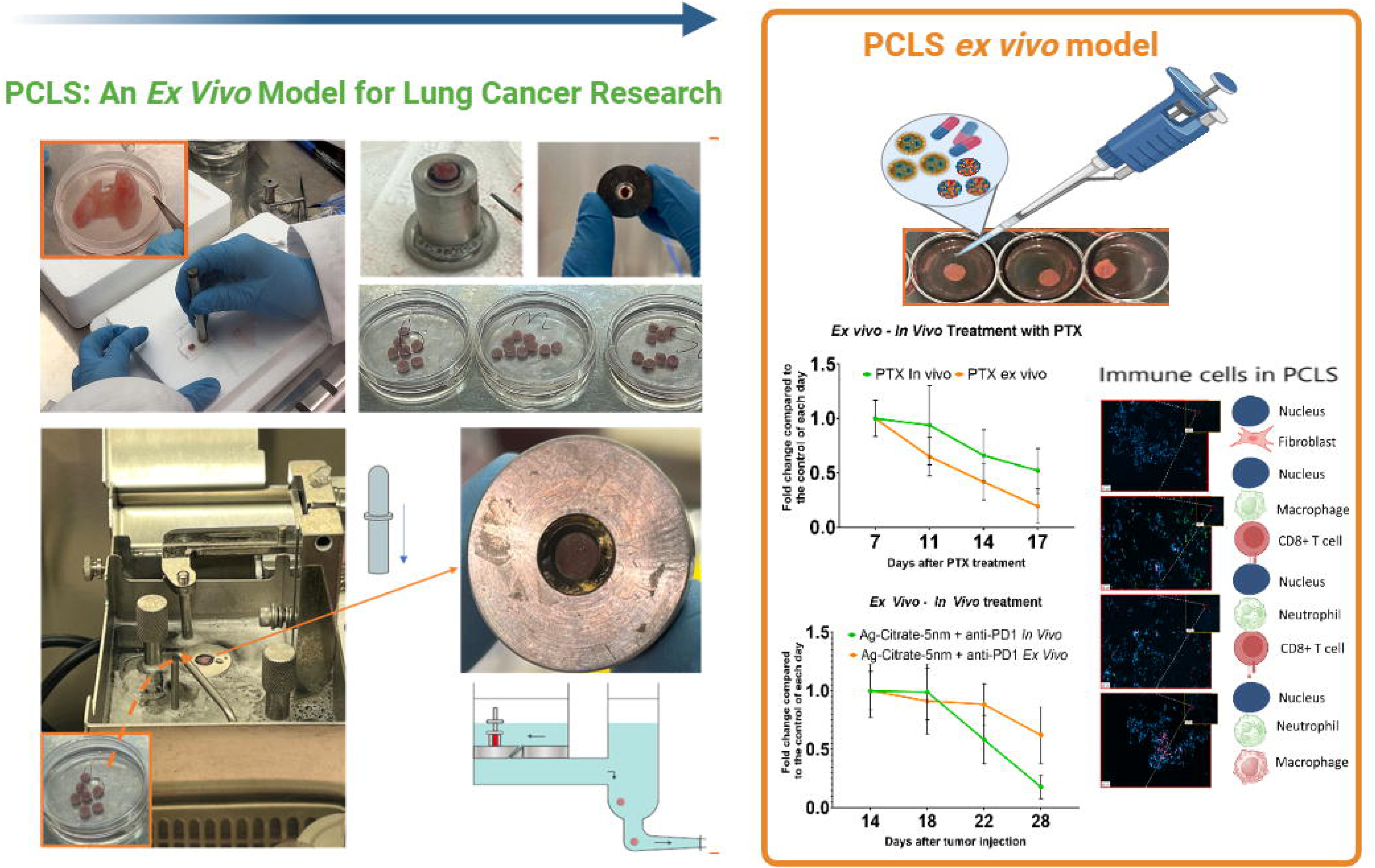

## Notes

### Competing Interest Statement

The authors have declared no competing interest.

